# Inter-species gene flow drives ongoing evolution of *Streptococcus pyogenes* and *Streptococcus dysgalactiae* subsp. *equisimilis*

**DOI:** 10.1101/2023.08.10.552873

**Authors:** Ouli Xie, Jacqueline M. Morris, Andrew J. Hayes, Rebecca J. Towers, Magnus G. Jespersen, John A. Lees, Nouri L. Ben Zakour, Olga Berking, Sarah L. Baines, Glen P. Carter, Gerry Tonkin-Hill, Layla Schrieber, Liam McIntyre, Jake A. Lacey, Taylah B. James, Kadaba S. Sriprakash, Scott A. Beatson, Tadao Hasegawa, Phil Giffard, Andrew C. Steer, Michael R. Batzloff, Bernie W. Beall, Marcos D. Pinho, Mario Ramirez, Debra E. Bessen, Gordon Dougan, Stephen D. Bentley, Mark J. Walker, Bart J. Currie, Steven Y. C. Tong, David J. McMillan, Mark R. Davies

**Affiliations:** Department of Infectious Diseases, the University of Melbourne at the Peter Doherty Institute for Infection and Immunity, Melbourne, Australia; Monash Infectious Diseases, Monash Health, Melbourne, Australia; Department of Microbiology and Immunology, the University of Melbourne at the Peter Doherty Institute for Infection and Immunity, Melbourne, Australia; Menzies School of Health Research, Charles Darwin University, Darwin, Australia; European Molecular Biology Laboratory, European Bioinformatics Institute EMBL-EBI, Hinxton, UK; Australian Infectious Diseases Research Centre and School of Chemistry and Molecular Biosciences, The University of Queensland, Brisbane, Australia; Doherty Applied Microbial Genomics, Department of Microbiology and Immunology, The University of Melbourne at The Doherty Institute for Infection and Immunity, Melbourne, Australia; Department of Biostatistics, University of Oslo, Blindern, Norway; Faculty of Veterinary Science, The University of Sydney, Australia; Infection and Inflammation Laboratory, Queensland Institute of Medical Research Berghofer (QIMR) Berghofer Medical Research Institute, Brisbane, Australia; School of Science & Technology, University of New England, Armidale, NSW, Australia; Department of Bacteriology, Nagoya City University Graduate School of Medical Sciences, Nagoya, Japan; Tropical Diseases, Murdoch Children’s Research Institute, Parkville, Australia; Institute for Glycomics, Griffith University, Southport, Australia; Respiratory Disease Branch, National Center for Immunizations and Respiratory Diseases, Centers for Disease Control and Prevention, Atlanta, Georgia, USA; Instituto de Microbiologia, Instituto de Medicina Molecular, Faculdade de Medicina, Universidade de Lisboa, Lisboa, Portugal; Department of Pathology, Microbiology and Immunology, New York Medical College, Valhalla, NY, USA; Sanger Institute, Wellcome Trust Genome Campus, Hinxton, Cambridge, UK; Institute for Molecular Bioscience, The University of Queensland, Brisbane, Australia; Victorian Infectious Disease Service, The Royal Melbourne Hospital, Peter Doherty Institute for Infection and Immunity, Melbourne, Victoria, Australia; School of Science and Technology, Engineering and Genecology Research Centre, University of the Sunshine Coast, Maroochydore, QLD, Australia

**Keywords:** Streptococcus dysgalactiae, Streptococcus pyogenes, gene flow, population genomics

## Abstract

*Streptococcus dysgalactiae* subsp. *equisimilis* (SDSE) is an emerging cause of human infection with invasive disease incidence and clinical manifestations comparable to the closely related species, *Streptococcus pyogenes*. Through systematic genomic analyses of 501 disseminated SDSE strains, we demonstrate extensive overlap between the genomes of SDSE and *S. pyogenes.* More than 75% of core genes are shared between the two species with one third demonstrating evidence of cross-species recombination. Twenty-five percent of mobile genetic element (MGE) clusters and 16 of 55 SDSE MGE insertion regions were found across species. Assessing potential cross-protection from leading *S. pyogenes* vaccine candidates on SDSE, 12/34 preclinical vaccine antigen genes were shown to be present in >99% of isolates of both species. Relevant to possible vaccine evasion, six vaccine candidate genes demonstrated evidence of inter-species recombination. These findings demonstrate previously unappreciated levels of genomic overlap between these closely related pathogens with implications for streptococcal pathobiology, disease surveillance and prevention.

## Introduction

*Streptococcus dysgalactiae* subspecies *equisimilis* (SDSE), a beta-hemolytic *Streptococcus* normally possessing the Lancefield group C/G antigen, is an emerging cause of human disease with recently reported incidences of invasive disease comparable to or surpassing that of the closely related and historically important pathogen, *Streptococcus pyogenes* (group A *Streptococcus*) ^1–8^. SDSE and *S. pyogenes* occupy the same ecological niche and possess overlapping disease manifestations including pharyngitis, skin and soft tissue infections, necrotising fasciitis, streptococcal toxic shock syndrome and osteoarticular infections^9, 10^.

Gene transfer between SDSE and *S. pyogenes,* including housekeeping multi-locus sequence typing (MLST) loci, major virulence factors including the *emm* gene, and antimicrobial resistance (AMR) determinants has been reported^11–17^. Inter-species transfer of genes encoding the serogroup carbohydrate have led to SDSE isolates which express the group A carbohydrate^15, 18^. Transfer of accessory virulence or AMR genes is thought to occur by cross-species exchange of mobile genetic elements (MGEs) such as prophage or integrative conjugative elements (ICE) ^19–21^. However, the mechanism that enables the exchange of genes present in all strains of a species, termed ‘core’ genes, is not well understood. The extent of genetic exchange between these two pathogens has not been defined within a global population genomic framework.

An analysis of the population structure of a globally diverse collection of *S. pyogenes* genomes provided insights into the drivers of population diversity and global utility of candidate vaccines. In contrast, studies of SDSE whole genome diversity have generally been limited to local jurisdictions^7, 22–25^. With increasing efforts to develop a *S. pyogenes* vaccine^26, 27^, an improved understanding of the overlap and extent of genetic similarity between human isolates of SDSE and *S. pyogenes* in a global context is required. Here, we have compiled a globally diverse collection of 501 SDSE genomes isolated from human hosts. Through a gene synteny-based approach, we conducted a systematic analysis of the population genetics and pangenome of SDSE. Using the same framework, we compare SDSE to a previously published global *S. pyogenes* dataset^22^ to reveal extensive genomic overlap between the two closely related pathogens including genes encoding candidate *S. pyogenes* vaccine antigens.

## Results

### A globally diverse SDSE genomic database

To assess SDSE population diversity (Figure 1), we compiled a genomic database of 501 geographically distributed SDSE isolates from 17 countries including 228 newly reported genomes with one new complete reference quality genome, NS3396^28^ (Supplementary Table 1a). The database includes 53 *emm* sequence types, 88 *emm* sub-types, and 129 MLSTs.

**Figure 1.**
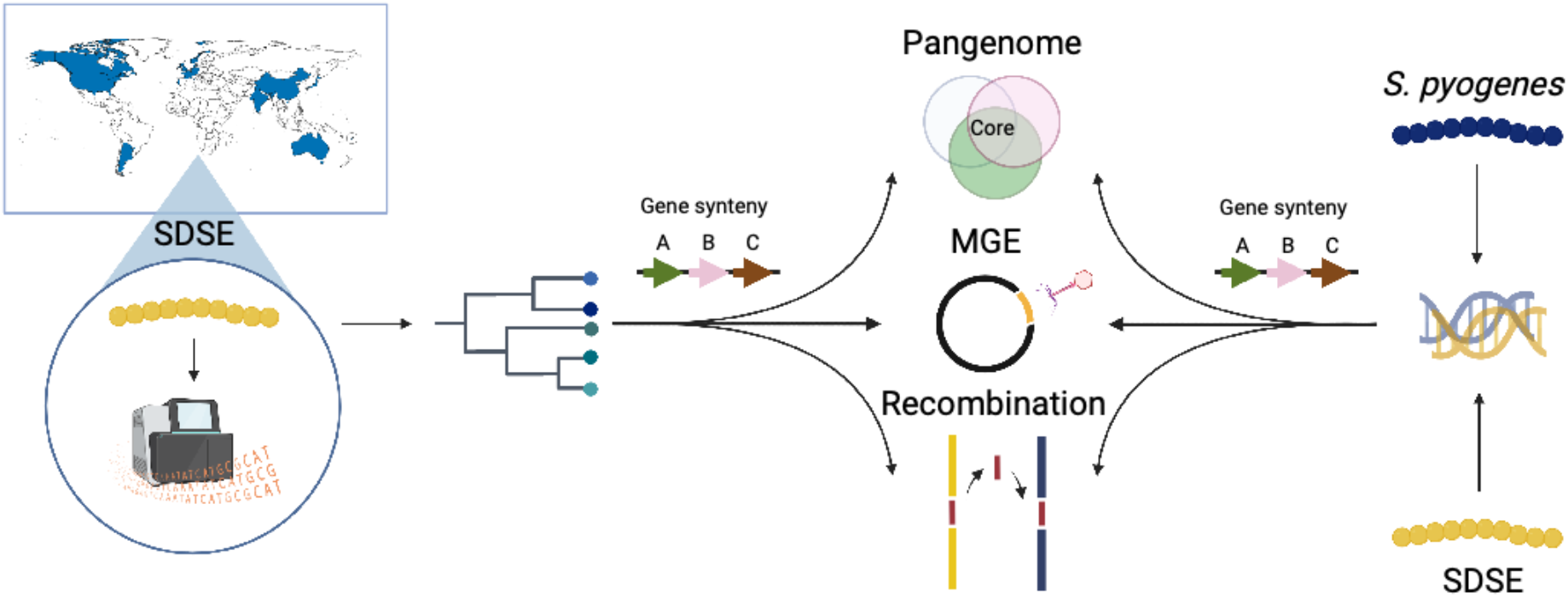
Workflow to characterise the global population structure of *Streptococcus dysgalactiae* subsp. *equisimilis* (SDSE) and its overlap with *S. pyogenes*. A globally diverse collection of 228 SDSE isolates were whole genome sequenced using Illumina short read sequencing and collated with publicly available genomes to form a database of 501 global sequences. An analysis of the SDSE population structure was undertaken followed by a systematic analysis of the pangenome, evidence of homologous recombination, mobile genetic elements (MGEs) and MGE insertion sites using pangenome gene synteny contextual information. This framework was then compared with a previously published global database of 2,083 *S. pyogenes* genomes by merging pangenomes accounting for shared gene synteny to reveal extensive overlap at the level of shared genes, homologous recombination and MGEs^22^. Figure created with BioRender.com.

### Global SDSE population structure

A phylogeny of the global SDSE population was constructed using a recombination aware pipeline implemented in Verticall. The minimum evolution phylogeny demonstrated a deep radially branching structure forming multiple distinct lineages similar to that observed for the global population structure of *S. pyogenes*^22^ (Figure 1a and Supplementary Figure 1).

Whole genome clustering of the global SDSE population using PopPUNK^29^ detected 59 distinct population clusters (akin to ‘lineages’ or ‘genome clusters’) (Figure 2a and Supplementary Figure 2). These genome clusters were geographically dispersed and were highly concordant with the inferred phylogeny.

**Figure 2.**
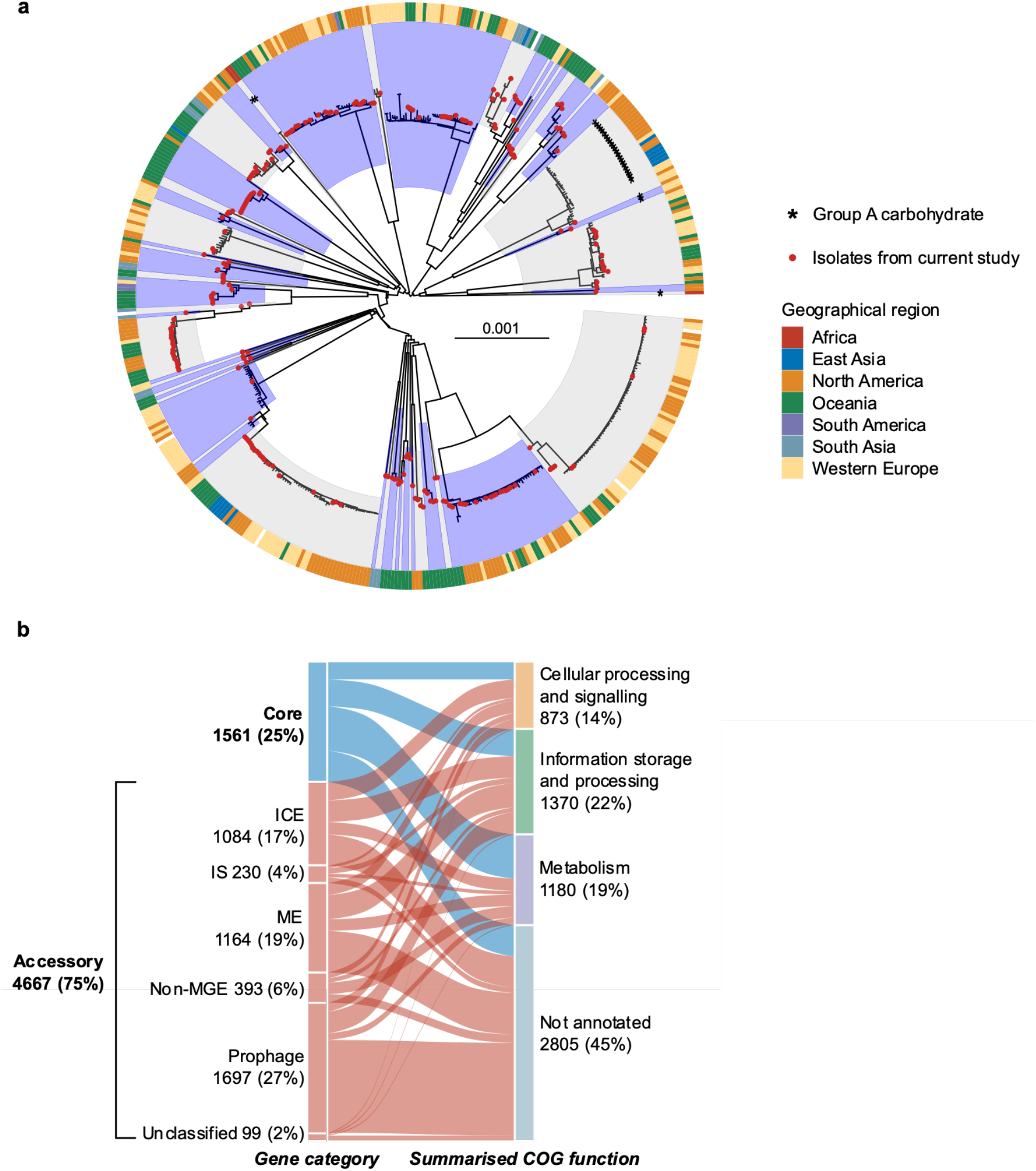
Global population structure and pangenome composition of *Streptococcus dysgalactiae* subsp. *equisimilis* (SDSE). **a)** Minimum evolution phylogenetic tree of 501 SDSE isolates using recombination masked genomic distances. Fifty-nine distinct whole genome clusters are highlighted by alternating blue and grey shades from internal nodes. Newly published sequences from this study are highlighted by red points at the tips of the tree. Isolates carrying the group A carbohydrate are marked by asterisks. The outer ring around the tree denotes geographical region of isolation. **b)** Alluvial plot of the SDSE pangenome with categorisation of core (present in ≥99% of isolates) and accessory genes into summarised COG functional groups. Accessory genes are further characterised based on the type of element they are most frequently found. Genes associated with prophage and prophage-like elements were grouped together. Genes associated with mobile genetic elements (MGE) that could not be classified further were assigned to mobility elements (ME). A small number of unclassified accessory genes were present only adjacent to contig breaks without an integrase and could not confidently be classified into a MGE or non-MGE category. ICE, integrative conjugative element; IS, insertion sequence; non-MGE, non-mobile genetic element.

Our data show a limited concordance between the inferred whole genome population structure and the classical SDSE molecular epidemiological markers, *emm* type and MLST (Supplementary Figure 3). Of the 24 *emm* types represented by three or more isolates, 11 (46%) were present in multiple genome clusters. Of the 27 genome clusters with three or more isolates, 18 (67%) contained two or more *emm* types.

MLST was in stronger agreement with the inferred genome clusters than *emm* type. Nevertheless, the largest genetic distance between isolates in the same MLST was frequently greater than that separating isolates from different MLST types (Supplementary Figure 3d). Supporting this observation, 67% (18/27) of genome clusters with three or more members contained more than one MLST. While many of these differed at only a single MLST locus, single locus variant MLST clonal complexes delineated distant lineages poorly and were present across multiple distinct genetic backgrounds (Supplementary Figure 3e). These findings indicate that while *emm* and MLST typing for SDSE have been useful epidemiological markers in jurisdictional or regional contexts, they have limitations when defining global SDSE evolution and diversity. The presented whole genome clustering scheme implemented in PopPUNK provides a solution to the limitations of *emm* and MLST at the global population level^29^.

The Lancefield group carbohydrate was predicted by the presence of a 14 gene carbohydrate synthesis gene cluster in group C SDSE^18^, 15 gene cluster in group G^15^ and 12 gene cluster in group A *Streptococcus*^15^ (Supplementary Figure 4a) in a conserved genomic location immediately upstream of *pepT*. Of the 27 genome clusters which contained three or more isolates, 16 consisted of only group G isolates and 9 consisted of only group C (Supplementary Figure 4b). Two genome clusters consisted of isolates with more than one group carbohydrate. The Lancefield group A carbohydrate, normally associated with *S. pyogenes*, was found in isolates from 4 different genome clusters (Figure 2a), including one genome cluster that also contained isolates with the group C and group G antigen. These findings suggest that while closely related isolates generally express the same Lancefield group carbohydrate antigen, horizontal transfer of the group carbohydrate locus can occur, including the group A carbohydrate locus normally associated with *S. pyogenes*.

Bayesian dating of the most recent common ancestor (MRCA) of the two most well sampled genome clusters using BactDating^30^ with recombination masked by Gubbins^31^, revealed emergence of these lineages within the last 100 years. A genome cluster containing members present in 7 countries and inclusive of *emm* type *stG*6792/ST 17 isolates, until recently the most frequent cause of invasive disease in Japan^7, 10, 32^, had an estimated root date in the mid-1970s (credible interval CrI 1960 – 1991, Supplementary Figure 5a) suggesting recent emergence and global dispersion. A genome cluster consisting of isolates with *emm* type *stG*62647, an *emm* type described to be increasing in frequency in multiple jurisdictions^24, 33^, had an estimated root date in the early 1930s (CrI 1855 – 1969, Supplementary Figure 5b). These findings suggest that major genome clusters in this global database represent modern, globally disseminated lineages.

Virulence genes or regulators known to influence expression of virulence genes were enriched in a subset of genome clusters. Isolates were screened for the presence of accessory toxins including DNAses, superantigens, adhesins, immune escape factors and regulators described in SDSE or *S. pyogenes* (Supplementary Table 1b). We found that certain genome clusters were enriched for accessory factors including: chromosomally encoded exotoxin *speG*, immune escape factor *drsG*, adhesin *gfbA* and an accessory/secondary FCT locus, presence of the negative regulatory *sil* locus, and three accessory prophage streptodornase genes *spd3*, *sda1* and *sdn* (all p <0.001, χ^2^ test of independence). In particular, the accessory FCT locus^34^, *sil* locus, *gfbA*, *speG* and *drsG* were frequently either present or entirely absent from different genome clusters. These findings suggest that the SDSE genetic population structure is characterised by distinct virulence repertoires. Non-random sampling prevented associations with clinical manifestations from being inferred.

### Signatures of recombination in the SDSE core genome

The SDSE pangenome consisted of 6,228 genes of which 1,561 were core genes (present in ≥99% of genomes) and 4,667 were accessory genes, of which 3,849 genes were present in <15% of genomes (Figure 2b). While several SDSE genes have previously been reported to be recombinogenic, we next aimed to infer the number of SDSE core genes with signatures of recombination in their evolutionary history using fastGEAR^35^. Of 1,543 core genes (excluding 18 genes that had over 25% gaps in their alignment), 837 (54%) genes had a recombinatorial signature (Supplementary Table 2a). These notably included all seven MLST genes. These findings correlate with uncertainties when classifying isolates using MLST compared to whole genome clusters.

To further quantify the contribution of recombination to SDSE population diversity, we measured the ratio of recombination-derived mutation vs vertically inherited mutation (r/m) using Gubbins^31^ for the 12 largest genome clusters (385 isolates). The median r/m per genome cluster was 4.46 (range 0.38 – 7.05) which was comparable to a r/m of 4.95 estimated previously for *S. pyogenes*^22^ (Supplementary Table 3). The median recombination segment length was 4,647 bp (range 6 – 97,789 bp) and the median number of events per genome cluster was 42 (range 3 – 64), again similar to that previously described for *S. pyogenes*^22^, supporting multiple small fragments of homologous recombination as a major source of genetic diversity in SDSE.

### Contribution of the accessory genome to SDSE diversity

To investigate the contribution of MGE and non-MGE genes to accessory genome diversity, an identification and characterisation workflow was developed using genome synteny and a classification algorithm adapted from proMGE^36^. Segments of accessory genes (hereafter referred to as ‘accessory segments’) were classified as prophage, phage-like, ICE, insertion sequence/transposon (IS) or non-MGE based on the presence of MGE-specific integrase/recombinase genes and prophage or ICE structural genes (Supplementary Figure 6a). Accessory segments were classed as mobility elements (ME) when insufficient information was present to classify a segment – such as with degraded elements, complex nested elements or elements fragmented by assembly breaks with insufficient contextual information. Genes were classified into MGE categories based on the frequency of their presence in different elements (Figure 2b). Prophage elements contributed the largest number of genes to the accessory genome (36%, 1,697/4,667 genes) followed closely by ME (25%, 1,164/4,667 genes) and ICE (23%, 1,084/4,667 genes). Non-MGE accessory genes constituted 8% (393/4,667 genes) of the accessory genome. A mean of 0.8 prophage (range 0 – 5), 1.1 phage-like (range 0 – 4), 1.1 ICE (range 0 – 4) and 4.7 ME (range 0 – 10) were found per genome. MGE counts were similar when restricted to four complete SDSE genomes (mean 1.3 prophage, 1 phage-like, 1 ICE, 2.8 ME). A mean of 10.5 IS/transposon elements (range 3 – 30) were found per genome but was likely to be an underestimation as parameters used to construct the pangenome resulted in the removal of infrequent genes at genome assembly breaks, which are commonly associated with IS/transposons.

Using genome location defined between two syntenic core genes, MGE chromosomal ‘insertion regions’ were mapped in the SDSE dataset. A total of 55 insertion regions were found in SDSE (Figure 3a, Supplementary Table 4a). Prophage or phage-like elements were found at 32 insertion regions (58%), ICE at 21 regions (38%), and 10 regions (18%) were occupied by either prophage or ICE. Twelve regions (22%) were occupied by elements classified as ME only, reflective of the limitations imposed by sequence breaks around MGEs in draft genomes. The MGE insertion regions were broadly occupied across SDSE genome clusters (Supplementary Figure 6b).

**Figure 3.**
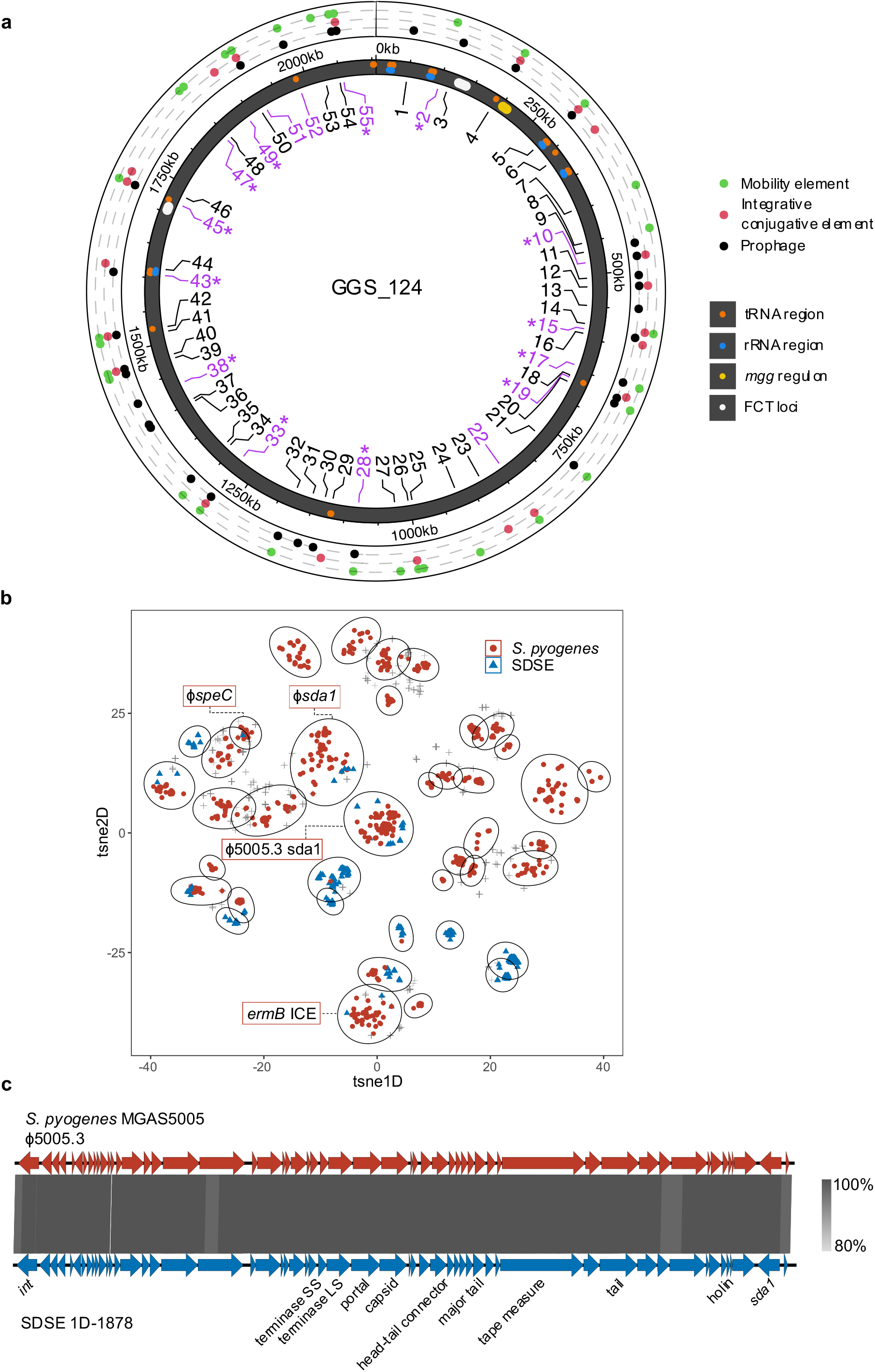
Clustering and genome localisation of *Streptococcus dysgalactiae* subp. *equisimilis* (SDSE) and *S. pyogenes* mobile genetic elements (MGEs). **a)** Location of SDSE MGE insertion regions relative to the GGS_124 reference genome (NC_012891.1). Insertion regions are labelled from 1 to 55 of which 16 (29%) were shared with *S. pyogenes* (highlighted in purple). Shared insertion regions at which MGE clusters were present and shared across species are highlighted with asterisks. The outer ring indicates the type of element detected at each insertion region in SDSE. tRNA and rRNA regions, the *mgg* regulon containing the *emm* gene, and the two FCT loci in GGS_124 are marked on the genome representation. **b)** Clustering of 3,630 MGEs at 16 shared insertion regions across SDSE and *S. pyogenes*. Each cluster is outlined by an ellipsoid. Of the 40 MGE clusters, 10 (25%) were shared across species. Clusters which contained notable examples of near-identical (>80% nucleotide identity and coverage) cross-species MGEs are labelled: the global M1T1 ϕ5005.3 prophage carrying the Sda1 streptodornase (104 *S. pyogenes*, 12 SDSE), ϕ*sda1* refers to a prophage carrying a *sda1* allele 95% similar to ϕ5005.3 (2 *S. pyogenes*, 1 SDSE), ϕ*speC* refers to a previously described prophage carrying *speC* and *spd1*^18^ (2 *S. pyogenes*, 1 SDSE), *ermB* ICE refers to a complex nested ICE and IS/transposon element carrying multiple AMR genes including *ermB* (1 *S. pyogenes*, 1 SDSE). These MGEs represent a subset of elements within each respective cluster. **c)** Genome architecture and comparison of the *S. pyogenes* M1T1 prophage ϕ5005.3, with a near identical prophage found in group G SDSE isolate 1D-1878 at insertion site 45. 1D-1878 was isolated from a case of invasive disease in Denmark in 2018^53^. Regions of genomic similarity were inferred using BLAST and plotted using Easyfig v2.2.3^76^. The grey gradient indicates the percent identity in the legend. The same prophage was detected in an additional 19 SDSE isolates spanning 10 distinct genome clusters and 7 countries indicating significant dispersion of the prophage in the SDSE population.

### Conservation of MGE insertion regions across SDSE and S. pyogenes are associated with shared elements

Applying the same accessory identification and categorisation workflow to the 2,083 *S. pyogenes* genomes published previously^22^ enabled a systematic comparison of MGE and associated insertion sites both within and between the two species. An average of 1.9 prophage (range 0 – 6), 0.4 phage-like (range 0 – 3), 0.2 ICE (range 0 – 2), and 1.9 ME (range 0 – 7) were found per *S. pyogenes* genome. Compared to SDSE, *S. pyogenes* isolates had more prophage elements but less ICE (p < 0.001, Wilcoxon rank sum).

Overlaying the SDSE and *S. pyogenes* pangenomes while accounting for genome synteny, 31 previously published^37, 38^ and 13 new *S. pyogenes* MGE insertion regions (31 prophage, 9 ICE, and 2 mixed prophage and ICE) were mapped and compared to 55 insertion regions in SDSE (Supplementary Table 4c). Of these, 16 (29% of SDSE insertion regions) insertion regions were shared across the species (Figure 3a). At the 16 cross-species insertion regions, 1,443 accessory genes (54% of SDSE and 59% of *S. pyogenes* accessory genes at these regions) were shared across species, suggesting likely shared MGEs at these regions. At insertion regions which were not conserved across the two species, 816 accessory genes (34% of SDSE and 42% of *S. pyogenes* accessory genes) were shared, significantly less than the proportion at conserved insertion regions (p < 0.001, χ^2^ test).

Hypothesising that shared MGE insertion regions may provide a basis for shared prophage and ICE between species, all completely assembled and fragmented putative MGEs >10-15kb at the 16 cross-species insertion regions were examined. A total of 3,335 fully assembled and 295 high-quality fragmented putative MGEs, 710 from SDSE and 2,920 from *S. pyogenes*, were extracted. To identify similar elements within this database, we used mge-cluster^39^ to cluster ICE, ME and prophage elements which overcomes limitations of sequence homology-based approaches which are restricted by the modular and highly recombinogenic nature of MGEs. Using this approach, 40 clusters containing 2,897 MGEs were identified of which 10 (25%) ICE and prophage clusters across 13 insertion regions were found in both species (Figure 3b). The clusters were generally well defined and separated by MGE type (Supplementary Figure 7). Of the 40 clusters, 14 were present at more than one insertion region (median 1, range 1 – 7) indicating broad sharing of MGE clusters across these common insertion regions and between species.

We further examined MGE clusters for examples of shared near-identical elements (at least >89% nucleotide identity and coverage). Four MGEs (prophage, ICE and nested IS/transposon elements) carrying streptodornase genes *sda1* and *spd1*, the exotoxin gene *speC*, and AMR genes *ermB*, *ant(6)-Ia* and *aph(3’)-IIIa* were detected across both species (Table 1, Supplementary Figure 8). A completely assembled prophage element that was near-identical (>99% nucleotide identity) to the *S. pyogenes* M1T1 prophage ϕ5005.3 was shared across 12 genomes in SDSE and 104 genomes in *S. pyogenes* (Figure 3c). The M1 prophage ϕ5005.3 carries the streptodornase gene *sda1*, an extracellular virulence factor thought to have been acquired during the emergence of the global M1T1 *S. pyogenes* lineage^40, 41^. To investigate the distribution of this prophage further in SDSE, including MGEs fragmented by contig breaks, co-presence of the same integrase and *sda1* allele was examined across the SDSE database. An additional 8 genomes with co-carriage of these genes at the same insertion region were found. The ϕ5005.3 prophage was inferred to be present in SDSE isolates spanning 10 different genome clusters across seven countries (Australia, Canada, China, Denmark, Norway, Portugal and USA), over a 20-year period (1999 to 2018). Within these genome clusters, a mean of 28.8% of isolates carried the ϕ5005.3 phage (range 1.5 – 100%). These findings indicate that MGEs carrying virulence and/or AMR determinants can cross species boundaries including evidence of dispersal into more than one global SDSE lineage.

**Table 1.**
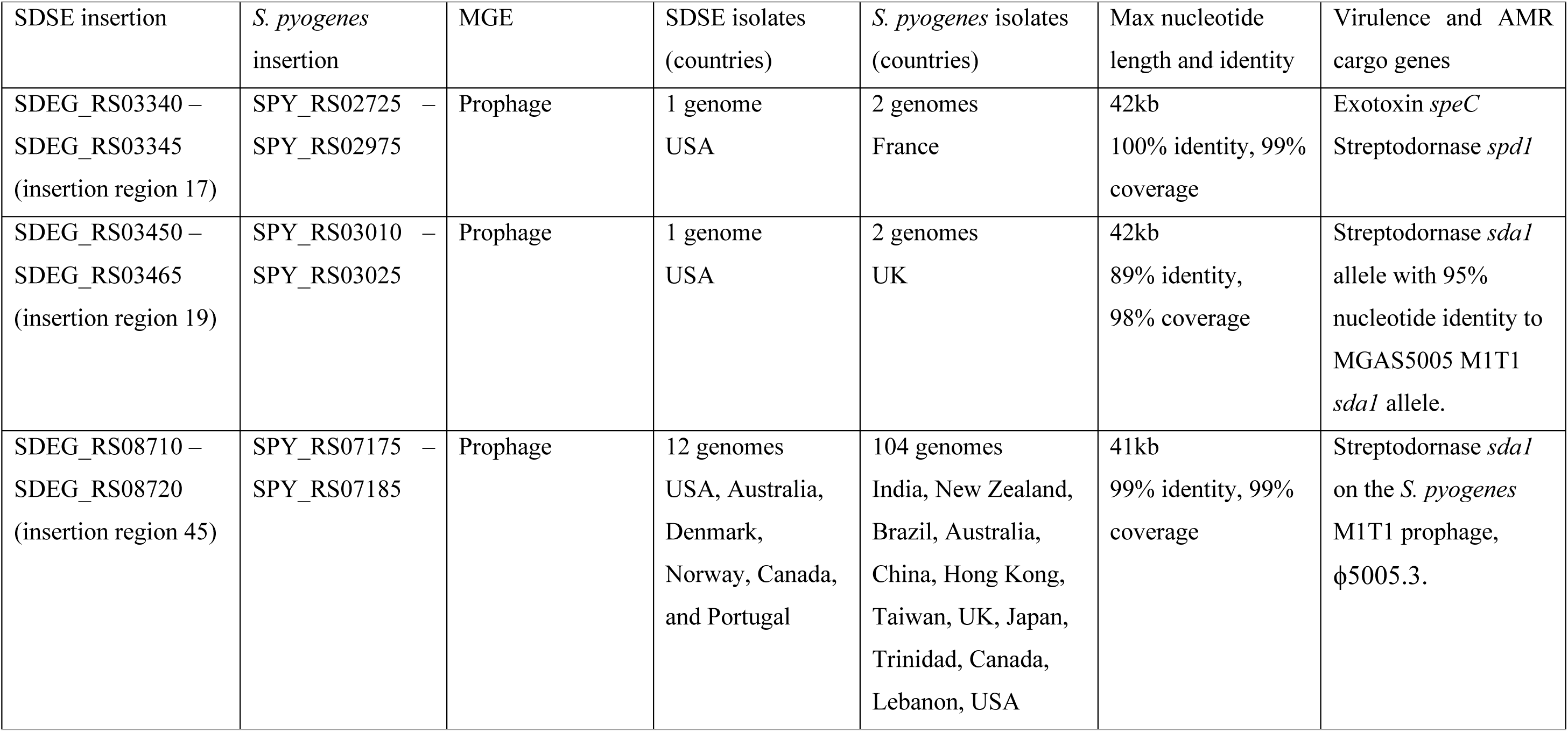
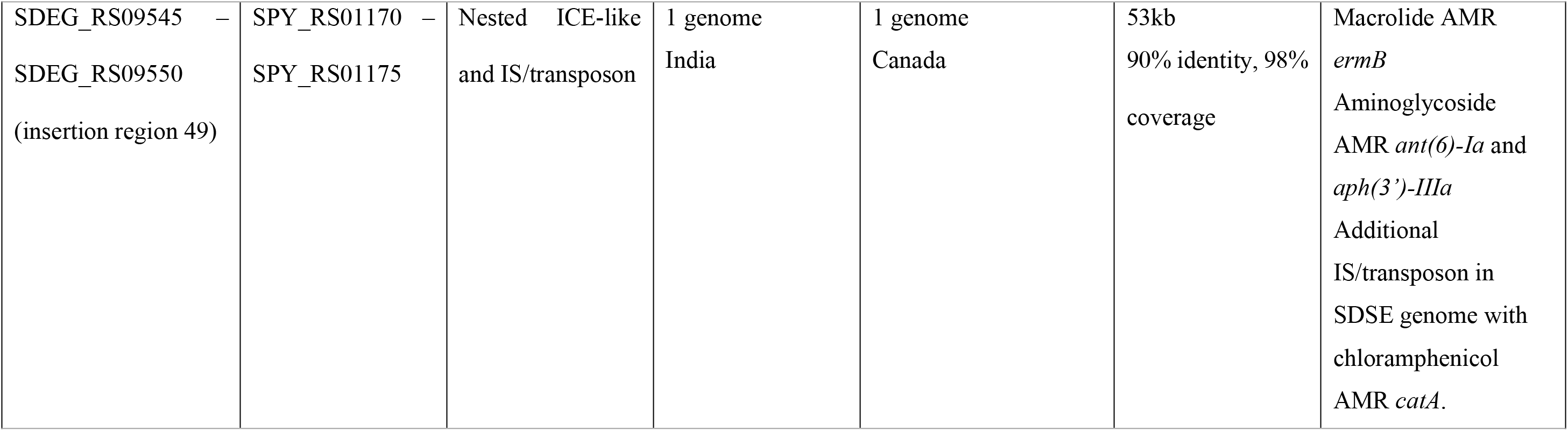
Fully assembled mobile genetic elements (MGE) shared at conserved cross-species insertion regions in the global *Streptococcus dysgalactiae* subsp. *equisimilis* (SDSE) and *S. pyogenes* databases. SDSE insertion sites are mapped to reference genome GGS_124 (NC_012891.1) and *S. pyogenes* insertion sites are mapped to reference genome SF370 (NC_002737.2). SDSE insertion region number refers to naming in Figure 3a. Antimicrobial resistance, AMR; ICE, integrative conjugative element; IS, insertion sequence.

### Extensive overlap and exchange in the core genome between SDSE and S. pyogenes

To examine core genome overlap across SDSE and *S. pyogenes*, we found that 1,166 core genes comprising 75% (1,166/1,547) of the SDSE core genome and 88% (1,166/1,320) of the *S. pyogenes* core genome were shared in the merged pangenome (Figure 4a, Supplementary Table 2b). A small number of genes/coding sequences were combined when merging pangenomes, resulting in a slightly smaller merged core pangenome compared to SDSE or *S. pyogenes* alone. To investigate cross species recombination in these core genes, 1,166 genes that were core in both species and with <25% length variation were aligned and assessed using fastGEAR^35^. A total of 526 core genes (45%) had clusters with members from both species consistent with either shared ancestry or whole gene recombination. Putative cross-species recombination was identified in 393 (34%) unique genes, including the MLST genes *gki*, *murI*, and *recP* and the penicillin-binding protein-encoding gene *pbp1b*, which have previously been documented to be affected by inter-species recombination^11, 16^ (Supplementary Table 2b). Of these, 216 genes had a signature of multiple unique events between species. Recombination affected genes across all functional categories with no significant difference between classes (p = 0.26, χ^2^ test of independence). While net directionality and absolute frequency of events cannot be inferred using this data, predicted cross-species recombination events affected genes from across the genome with few hot-spot regions of higher density or restriction (Figure 4b).

**Figure 4.**
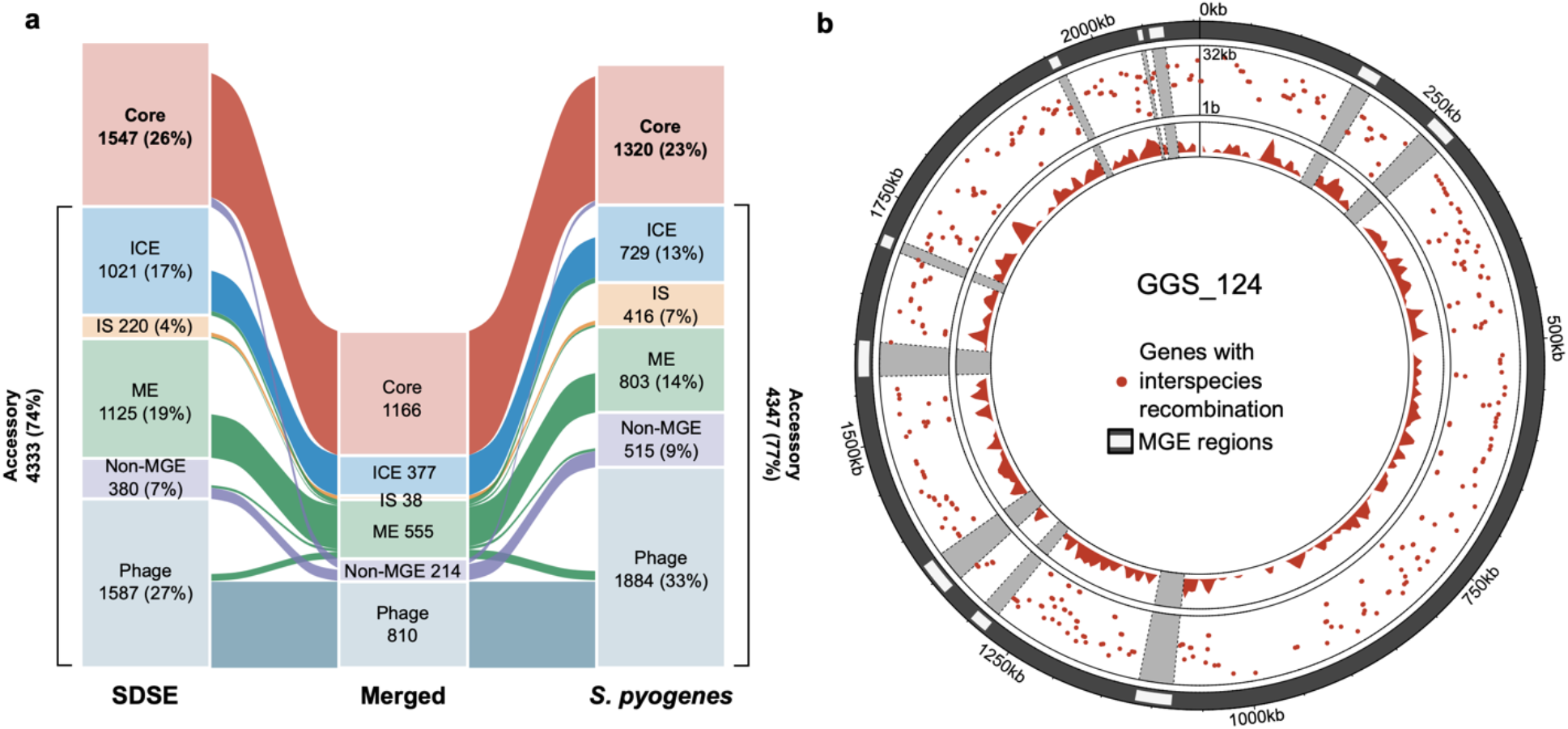
Comparison and recombination signatures between the *Streptococcus dysgalactiae* subsp. *equisimilis* (SDSE) and *S. pyogenes* pangenome. **a)** Alluvial plot of the shared SDSE (n=501) and *S. pyogenes* (n=2,083) pangenome. Categorisation of core genes (present in ≥99% of isolates in both species) and accessory genes by association with mobile genetic element (MGE). ICE, integrative conjugative element; IS, insertion sequence; ME, mobility element. Genes classified as belonging to different types of MGEs or an MGE/non-MGE combination across the species were binned as ME in the merged pangenome. Unclassified genes were excluded and a small number of genes were merged when overlaying pangenomes resulting in a slightly smaller merged pangenome than the pangenomes of individual species. **b)** Circular rainfall plot of core genes with signatures of recombination flagged by fastGEAR^35^ plotted relative to the GGS_124 reference genome (NC_012891.1). Genes flagged with putative interspecies recombination events are highlighted by red points. The distance between each gene with its neighbour within the same category is plotted on a log_10_ scale between 1 bp to 32 kbp. The innermost track plots the density of genes with evidence of interspecies recombination using a window size of 10,000 bp. MGE regions are masked in grey. The rainfall plot demonstrates that genes flagged as affected by inter-species recombinants are dispersed across the SDSE genome.

### Predicted conserved metabolic pathways between species identified by pangenome analysis

Examination of non-MGE genes unique to the pangenomes of each species found well-defined KEGG modules predicting metabolic differences between the species. Core to SDSE but absent from the *S. pyogenes* pangenome were modules encoding glycogen biosynthesis (M00854) and threonine biosynthesis (M00018). *S. pyogenes* is known to be auxotrophic for threonine and the absence of threonine and glycogen biosynthesis genes may reflect greater host dependence and/or adaptation. Unique to *S. pyogenes* were multiple genes encoding V/A-type ATPases (M00159). While additional differences are likely to exist beyond described KEGG modules, 69% of SDSE and 86% of *S. pyogenes* metabolic genes were shared indicating extensive overlap between the species (Supplementary Figure 9).

### Conservation of leading S. pyogenes vaccine candidates in the global SDSE population

Given the extensive overlap in gene content between SDSE and *S. pyogenes*, we next assessed the carriage of 34 leading *S. pyogenes* candidate vaccine antigens (Supplementary Table 5) ^22^ in the global SDSE population. Of the 26 candidate vaccine antigens using a full or near-full length gene product, 12 were highly represented in both species (present in >99% of isolates at 70% identity, Figure 5a). Mean amino acid sequence divergence in SDSE for these 12 candidates from the *S. pyogenes*-derived reference sequence varied from 80.9% to 99.9% (Supplementary Figure 10a). Of the four small peptide vaccine candidates, the multivalent N-terminal M protein and multivalent Tee antigen candidates, which were searched at 100% identity, only J8.0 (C-terminal fragment of M protein) was present in all SDSE isolates. However, none were present in >99% of isolates in both species (Figure 5b). Potential coverage by five of 11 leading preclinical multicomponent vaccines was >99% in both populations (Figure 5c, Supplementary Table 6).

**Figure 5.**
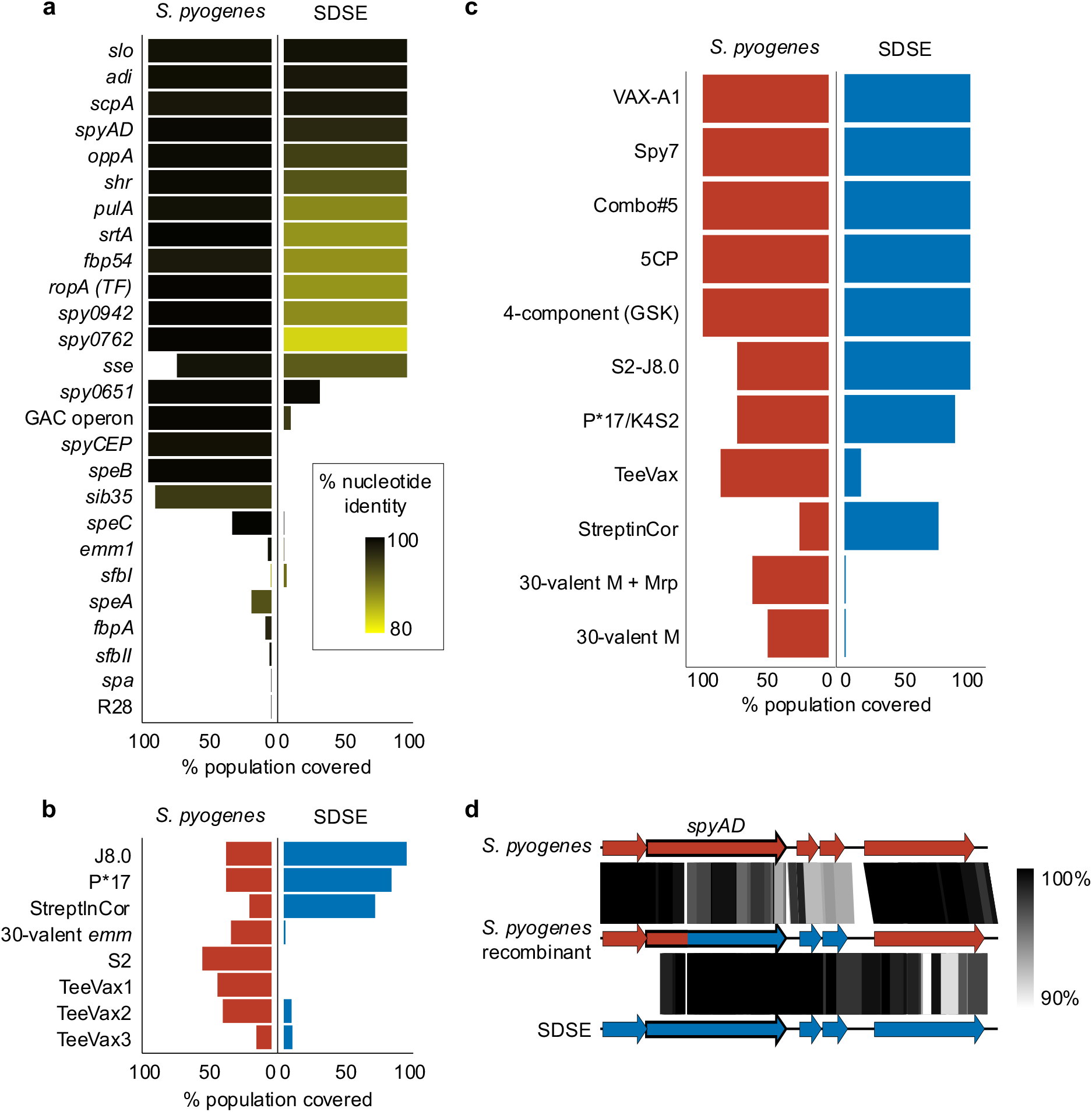
Theoretical coverage of preclinical vaccine candidates and multi-component formulations in global *Streptococcus dysgalactiae* subsp. *equisimilis* (SDSE, n = 501) and *S. pyogenes* (n = 2,083) populations. **a)** Whole gene candidates were screened in both species at 70% nucleotide length and identity. Theoretical coverage of the population (% presence) is expressed by length of bar and conservation relative to the *S. pyogenes* query sequence is reflected by gradient fill from 80% similar (yellow) to 100% (black). **b)** Peptides and gene sub-domain candidates were screened without calculation of sequence diversity. Peptide fragments were screened using a 100% match approach and a six-frame translation of the sequence (J8.0, S2, P*17, StreptInCor ‘common epitope’) and gene sub-domains (30-valent *emm*, T antigen fragments in TeeVax1, TeeVax2, TeeVax3) were screened at 70% nucleotide identity and coverage. **c)** Theoretical coverage of SDSE and *S. pyogenes* isolates in this study by multivalent vaccine candidates. **d)** Exemplar of putative recombination between *S. pyogenes* and SDSE vaccine candidate *spyAD*. *S. pyogenes* isolate NS1140 demonstrated evidence of cross-species recombination around the vaccine candidate *spyAD* gene. Sequence similarity between *S. pyogenes* reference genome SF370 NC_002737.2 (top), recombinant *S. pyogenes* isolate NS1140 (middle), and SDSE reference 3836-05 (bottom) demonstrated greater similarity between NS1140 and 3836-05 at position 419 of *spyAD* to 2 ORFs downstream. Regions of genomic similarity were inferred using BLAST and plotted using Easyfig v2.2.3^76^.

Shared antigens may provide cross-species vaccine coverage but conversely, interspecies recombination could yield increased antigenic diversity. Interspecies recombination analysis using fastGEAR was carried out on nine full length antigens which had <25% length variation and were highly present in both species. Five candidates, SpyAD, OppA, Shr, PulA and Fbp54, demonstrated signatures of recombination between a single-species cluster and a genome of the other species (Figure 5d and Supplementary Figure 10b). Three candidates, SLO, ADI and TF, had sequence clusters containing both species. Alleles of *srtA* were separated by species between SDSE and *S. pyogenes*. Examination of shared clusters found evidence of SDSE isolates which had acquired a TF allele from *S. pyogenes* (Supplementary Figure 10b). However, in the setting of limited sequence diversity, recombination could not be inferred for SLO and ADI.

## Discussion

With increasing disease control efforts for *S. pyogenes* including vaccine development^26, 27, 42^ and reports of high disease burden associated with invasive SDSE infection^1–8^, an improved understanding of the SDSE genomic population structure and overlap between these closely related pathogens provides a new framework for understanding the evolution and prevention of disease associated with infection by these human pathogens.

The population structure of SDSE was found to have many similarities to that of *S. pyogenes* with multiple evolutionary distinct lineages. Recombination of small genomic fragments was found to be a major driver of diversity of the core genome in SDSE with variation in the accessory genome due to MGE-related genes. These features mirror findings described previously for *S. pyogenes*, suggesting similar evolutionary dynamics at a global population level across these pathogens^22^.

At the level of the core genome, more than 75% of core genes were shared between each species. Despite the extensive overlap, metabolic differences were found in well characterised KEGG modules such as the presence of threonine and glycogen biosynthesis modules which are core in SDSE but absent in *S. pyogenes*. SDSE has on average, a larger genome than *S. pyogenes* (2.1Mbp vs 1.8Mbp). The threonine and glycogen biosynthesis modules were also present in 8 complete *Streptococcus dysgalactiae* subsp. *dysgalactiae* genomes available on RefSeq (accessed 6 March 2023). While theoretical, it is plausible that these metabolic differences may reflect greater levels of genome reduction in *S. pyogenes*, under the assumption that human isolates of SDSE may have more recently diverged from a multi-host reservoir^43^. We found extensive horizontal gene transfer through homologous recombination across the SDSE and *S. pyogenes* core genome, with over a third of SDSE genes demonstrating evidence of cross-species recombination. These findings extend previous recognition of such interspecies recombination of MLST genes^11, 17^ and is likely a combination of both ancestral and ongoing genetic transfers. While we do not quantify the frequency of cross-species recombination, the extent of recombination is comparable to that reported by Diop *et al.* who used measures of homoplasy and sliding window-based sequence identity^44^. This dataset may also provide a foundation for development of future methods controlling for population structure to infer net directionality and frequency of interspecies recombination.

Over 25% of MGE insertion regions and MGE clusters were found to be conserved across SDSE and *S. pyogenes*, suggesting much greater sharing of MGEs across the species than previously appreciated. Within MGE clusters, near-identical complete MGEs were detected including ϕ5005.3, the prophage carrying streptodornase Sda1 found in the globally successful M1T1 *S. pyogenes* lineage^41, 45^. The prophage was present across multiple distinct lineages in SDSE indicating sharing and dissemination of the element in the SDSE population.

Extending these findings to *S. pyogenes* vaccine candidates, 12 of 34 antigens and five of 11 multicomponent vaccine candidates were predicted to contain antigens present in >99% of isolates from both species. Of the 12 antigens, 6 had evidence of interspecies recombination including components of multivalent vaccines, SpyAD, OppA, TF and PulA. This suggests that SDSE may represent an additional reservoir for antigenic diversity particularly if vaccine candidates enhance selection in an antigenically diverse region. Even greater diversity likely exists considering current limited whole genome sampling of SDSE in the continents of Africa and South America. Thus, surveillance of SDSE should be considered in the context of *S. pyogenes* vaccine development and monitoring.

Although published data on preclinical efficacy of vaccine candidates in SDSE is limited beyond preliminary data on the J8 peptide^46^, the prevalence of conserved antigens suggest there may be cross-species effects of vaccines intended to target *S. pyogenes*. However, the sequence-based approach used in this study does not consider potential conserved structural epitopes which may confer antibody cross-reactivity or cross-opsonisation to divergent alleles. It should be noted though that an immunological correlate of protection has yet to be determined for *S. pyogenes* or SDSE.

This systematic and detailed analysis of the overlap between SDSE and *S. pyogenes* at the level of the core genome, recombination, and MGEs reveals the extensive shared genomic content between these closely related pathogens and provides a platform for further investigations into their shared biology. Genomic exchange however is not limited to movement between these two organisms. Indeed, horizontal gene transfer between SDSE and another beta-haemolytic *Streptococcus*, *Streptococcus agalactiae*, has been described and elements of SDSE biology such as a second FCT locus, are more closely associated with *S. agalactiae* than *S. pyogenes*^34^. These methods could therefore be applied to other closely related species to provide insight into the shared biology between disease-causing streptococci in humans.

## Methods

### Bacterial isolates

The collection of 501 global SDSE sequences included publicly available short-read sequence data from NCBI sequence read archive (SRA) and complete genome assemblies from NCBI RefSeq until 4 May 2022. These included studies of invasive and non-invasive SDSE from Japan^47–49^, Germany^50^, India^51^, China^52^, Canada^23^, Norway^24, 34, 43^, USA^18, 19^, Denmark^53^, Switzerland^54^ and The Gambia^55^. A further 228 invasive and non-invasive isolates were collected from Australia, France, USA, India, USA, Argentina, Trinidad, Japan, Fiji, and Portugal to maximise geographical diversity. Metadata for the isolates are available in Supplementary Table 1.

### Genome sequencing, assembly and quality control

For the newly sequenced genomes, microbial DNA was extracted and 75-100bp paired-end libraries were sequenced using the Illumina HiSeq 2500 platform (The Wellcome Sanger Institute, United Kingdom). Reads were trimmed using Trim Galore v0.6.6 (https://github.com/FelixKrueger/TrimGalore) with a Phred score threshold of 20-25 and filtered to remove reads <36bp. Reads were examined for contamination using Kraken2 v2.1.2^56^. Draft assemblies were generated using Shovill v1.1.0 (https://github.com/tseemann/shovill) with SPAdes assembler v3.14.0^57^ and a minimum contig length of 200bp. Only assemblies with <150 contigs (mean 94, range 57-149), total assembly size between 1.9-2.4Mb (mean, 2.11Mb, range 1.91 – 2.30Mb), and GC% between 38-40% (mean 39.3%, range 38.7 – 39.6%) were included. The mean N50 was 72,091bp (range 42,987 – 146,734bp). Annotations were generated using Prokka v1.14.6^58^ using the ‘–proteins’ flag with four Refseq *S. pyogenes* and SDSE genomes to supplement annotations for consistency. *emm* sequence typing was performed using emmtyper v0.2.0 (https://github.com/MDU-PHL/emmtyper) and MLST assigned using MLST v2.22.0 (https://github.com/tseemann/mlst)^59^. The Lancefield group carbohydrate was inferred by the presence genes in the 14 gene group C carbohydrate synthesis locus^18^, 15 gene group G^15^ locus and 12 gene group A^15^ locus in the SDSE pangenome. The carbohydrate synthesis locus was present in a conserved location between core genes *mscF* (SDEG_RS03715) and *pepT* (SDEG_RS03795). Reads for all newly sequenced genomes are available on SRA with accession numbers provided in Supplementary Table 1.

To make a complete genome sequence of the *S. dysgalactiae* subsp. *equisimilis* strain NS3396, high quality genomic DNA was extracted using the GenElute Bacterial Genomic DNA Kit (Sigma). The complete genome assembly of NS3396 (GenBank accession CP128987) was performed using SMRT analysis system v2.3.0.140936 (Pacific Biosciences). Raw sequence data was *de novo* assembled with the HGAP3 protocol. Polished contigs were error corrected with Quiver v1, and the final assembly structure check by mapping raw reads against the alignment with BridgeMapper v1 as previously described^60^.

Complete genomes were checked for genome arrangement and duplications around RNA operons using socru v2.2.4^61^. Genomes with unverified large-scale rearrangements and duplications around RNA operons compared to published complete genomes NC_012891.1 GGS_124, NC_018712.1 RE378 and NC_019042.1 AC-2713 were excluded. Four complete genomes were included in the final database.

### Phylogenetic analysis

Recombination masked distances between SDSE genomes were calculated using Verticall v0.4.0 (https://github.com/rrwick/Verticall) which conducts pairwise comparisons between genomes and masks regions with increased and/or decreased sequence divergence as putatively recombinogenic. Using recombination masked distances, a minimum evolution phylogenetic tree was generated using fastME v2.1.6.1^62^ with nearest neighbour interchange using BME criterion for optimisation and subtree pruning and regrafting.

A comparative maximum-likelihood tree, without masking of recombination, was generated by alignment of all genomes against reference genome NC_012891.1 GGS_124 using Snippy v4.6.0 (https://github.com/tseemann/snippy). MGE regions were masked and tree inference conducted with IQ-tree v2.0.6^63^ using a GTR+F+G4 model. Tree comparisons were performed using TreeDist v2.5.0^64^ to calculate generalised Robinson-Foulds distances with mutual clustering information and matched splits were visualised with the ‘VisualizeMatching’ function.

Bayesian dating of the MRCA of the two most sampled genome clusters (groups ‘1’ and ‘2’) was performed using BactDating v1.1^30^ with an additive relaxed clock model and 10^8^ iterations to ensure Markov Chain Monte Carlo convergence and parameter effective sample size >190. Recombination-masked alignments were used as input. Alignments were generated using Snippy v4.6.0 against a reference genome within the same genome cluster when available, NC_018712.1 for genome cluster ‘2’, or a high-quality draft genome (SRR3676046) for genome cluster ‘1’. Regions affected by recombination were masked using Gubbins v3.1.2^31^ and IQ-TREE with a GTR+G4 model for tree building, and set with maximum 10 iterations, minimum of 5 SNPs to identify a recombination block, minimum window size of 100bp and maximum window size of 10,000bp. Contigs were padded with 10,000 Ns, the maximum window size, for the genome cluster ‘1’ alignment to prevent Gubbins calling recombination across contigs. Four non-*stG*62647 isolates were excluded from genome cluster ‘1’ for dating analysis as they were >1500 SNPs distant and formed a distinct sub-lineage within the cluster.

### SDSE population genomics

Evolutionarily related clusters (genome clusters) were defined using PopPUNK v2.4.0^29^ which has previously been used to describe the population genomics of *S. pyogenes*. PopPUNK assigns clusters based on core and accessory distances calculated using sliding k-mers. Distances were calculated using k-mer sizes between 13 to 29 at steps of four (Supplementary Figure 3a). A three-distribution Bayesian Gaussian Mixture Model (BGMM) was fit with 2D cluster boundary refinement to obtain a network score of 0.93 and was chosen after benchmarking against a model built using HDBSCAN and BGMM models with different numbers of distributions (Supplementary Figure 3b and 3c). The PopPUNK model (v1) and genome cluster designations from this study are available at https://www.bacpop.org/poppunk/ and can be iteratively expanded.

Genomic distances used to compare PopPUNK genome clusters with traditional epidemiological markers, *emm* and MLST, were recombination masked distances calculated by Verticall as above. Distances are analogous to 100% minus average nucleotide identity. Single locus variant MLST clonal complexes were calculated using the goeBURST algorithm as implemented in PHYLOViZ v2.0^65^.

Known SDSE and *S. pyogenes* virulence factors were screened in the SDSE database using screen_assembly v1.2.7^22^ with sequence accessions as listed in Supplementary Table 1b at 70% nucleotide identity and 70% coverage. Virulence factors were supplemented using Abricate v1.0.1 (https://github.com/tseemann/abricate) with VFDB^66^ at 70% nucleotide identity and 70% coverage. Genes with length variation and assembly breaks due to large repeat regions, were screened using Magphi v1.0.1^67^ for conserved 5’ and 3’ sequences at a distance equal to the maximum known length of the gene. Antimicrobial resistance genes were screened using Abritamr v1.0.6 (https://github.com/MDU-PHL/abritamr), a wrapper for AMRfinder plus v3.10.18^68^, at default 90% identity and 50% coverage.

### SDSE pangenome and MGE analysis

The SDSE pangenome was defined using Panaroo v1.2.10^69^ which utilises a pangenome graph-based approach for pangenome clustering. Panaroo was run in ‘strict’ mode with initial clustering at 98% length and sequence identity followed by a family threshold of 70% to collapse syntenic gene families. Core genes were defined as genes present in ≥99% of genomes, shell accessory genes between 15%-99% of genomes, and cloud accessory genes in <15% of genomes. COG functional categories were assigned to core genes using eggNOG-mapper v2.1.7^70^ with default Diamond mode. COG categories J, K, L, A, B and Y were summarised as genes involved in ‘information storage and processing’, categories T, D, V, U, M, N, O, W and Z were summarised as genes involved in ‘cellular processing and signalling’, categories C, G, E, F, H, I, P and Q were summarised as genes involved in ‘metabolism’.

MGE detection and classification was performed by adapting an automated classification algorithm by Khedkar *et al*. ^36^ and enhanced by Corekaburra^71^. Corekaburra maps core gene consensus synteny from pangenomes and was used to find stretches of accessory genes or ‘accessory segments’ between core genes which are used as anchor points. Full details of the logic for classification of MGEs is as described by Khedkar *et al.* ^36^ and Hidden Markov Models (HMMs) are available at http://promge.embl.de/. Briefly, accessory segments were investigated for integrase/recombinase subfamilies using HMMER v3.3.2^72^ ‘hmmsearch’ with model-specific gathering thresholds. Representative translated protein sequences from the pangenome were used and resulted in almost identical results compared to searches using sequences from individual genomes. Integrase/recombinase subfamilies were mapped to specific MGE classes: prophage, ICE, IS/transposon and ME (or mobility islands). For subfamilies associated with more than one MGE class, additional information including presence of prophage or ICE structural genes was required. Prophage structural genes were annotated using eggNOG-mapper v2.1.7^70^. At least two prophage genes with an appropriate integrase/recombinase were required to classify an element as a prophage. ICE structural genes were classified using HMMs from TXSScan^73^ and ‘hmmsearch’ with an E-value threshold of 0.001. Unlike the original method by Khedkar *et al.*, we required the presence of at least one coupling protein and one T4SS ATPase with an appropriate integrase/recombinase for classification of an ICE to improve specificity. Elements with a prophage specific integrase/recombinase, but no prophage structural genes were classified as phage-like. Accessory segments without enough contextual information or which contained both prophage and ICE structural genes were classified as ME. ME may also represent degraded elements, integrative mobilizable elements (IME), or accessory segments fragmented by sequence breaks with a disconnect between the integrase/recombinase and structural genes. The boundaries of nested segments with more than one recombinase were not resolved and we did not define the exact *attR* and *attL* sites.

Classification of accessory genes into non-MGE or MGE classes was based on the frequency each gene was detected in each class. Accessory segments with prophage or ICE structural genes but no integrase or unclassified segments with a sequence break did not contribute to the count as they may represent part of a fragmented MGE. Genes were classified as MGE-related if it appeared on any MGE more frequently than it was found on a non-MGE element. A small number of genes only present adjacent to contig breaks without an accompanying integrase were considered unclassified as they could not be confidently grouped into an MGE or non-MGE category. For the purposes of gene classification, phage-like elements were grouped with prophage.

### SDSE recombination detection

To examine evidence of recombination within the core SDSE genome, fastGEAR^35^ was run on 1,543 core gene alignments from the 501 SDSE strains included in the study. Gene alignments were performed using MAFFT v7.505^74^. Alignments with greater than 25% gap characters were excluded. fastGEAR infers population structure for each alignment, allowing for the detection of lineages or clusters that have ‘ancestral’ and ‘recent’ recombination events between them. Default parameters were used with a minimum threshold of 4 bp applied for the recombination length.

The relative ratio of mutation due to recombination to vertically inherited mutation (r/m) was determined for the 12 most frequently sampled genome clusters (385 isolates) using Gubbins v3.1.2^31^ as described above for phylogenetic and BactDating analysis. Complete genomes within the same genome cluster were available and used for alignment for four genome clusters. The remaining genome clusters were aligned against a high-quality draft genome within the same genome cluster. MGE regions were masked in the alignment. The r/m, number of recombination events and length of recombination segments was calculated for each genome within a genome cluster by summing along branches from the Gubbins output. The median r/m was calculated for each genome cluster, and the median of these values was given as the species r/m.

### MGE insertion site mapping across SDSE and S. pyogenes

SDSE MGE insertion regions were defined between two core genes using Corekaburra^71^ to call pangenome synteny. Only prophage, ICE and ME insertion sites were mapped. MGE counts presented in Supplementary Table 4a were estimated by summarising fragmented MGEs on the same contig as a core gene, to the corresponding core gene pair insertion region. Phage-like elements were grouped with prophages. Alternative insertion sites were defined as the less commonly found connection between two core genes and represent genome rearrangements around MGEs.

Insertion regions found in *S. pyogenes* were collated from McShan *et al.* ^37^ and Berbel *et al.* ^38^ in addition to 13 newly reported insertion sites using Corekaburra. Insertion regions were mapped across species by merging the SDSE and *S. pyogenes* pangenome. The *S. pyogenes* pangenome was defined using Panaroo v1.2.10^69^ with the same parameters as that used for SDSE. The pangenomes of the two species were then merged using ‘panaroo-merge’ which overlays pangenome graphs. Merging was performed with an initial clustering threshold of 90% identity and 90% length followed by a threshold of 70% for collapsing syntenic genes. Insertion sites in each species were then matched using the merged pangenome. Where no match was obtained, including where genes were core in only one species, core genes within two genes either side of the insertion region were manually inspected for a match.

### Shared genes and MGEs at insertion sites

MGE genes at shared insertion sites were determined using Corekaburra^71^ by listing all accessory genes present between core gene insertion sites which were conserved across SDSE and *S. pyogenes*. Genes were considered shared across SDSE and *S. pyogenes* if they overlapped in the merged pangenome.

MGEs were clustered using mge-cluster^39^ which maps distances between MGEs using presence and absence of shared unitigs, projects the distances in two dimensions using t-SNE and clusters similar elements using HDBSCAN. mge-cluster has been used with plasmid sequences and we now extend it to use with prophage and ICE. mge-cluster was run with t-SNE perplexity of 75 and HDBSCAN minimum cluster size of 30. MGE sequences were extracted using Magphi v1.0.1^67^ with insertion site core genes used as seed sequences, maximum seed distance determined by the largest distance between the respective core pair calculated by Corekaburra, and ‘--print_breaks’ selected which attempts to extract sequences across contig breaks. When cross-species seed hits could not be obtained using nucleotide sequences, input of the ‘--protein_seed’ flag with translated protein sequences were used. To ensure only high quality MGE sequences were obtained, only sequences longer than 10-15kb were included. Fragmented MGEs were concatenated for input into mge-cluster. Some segments were unable to be extracted by Magphi as seed core genes were present on small contigs resulting in difficulty determining whether to extract sequences in the 5’ or 3’ direction.

Highly similar MGEs present in SDSE and *S. pyogenes* were found by sequence-based clustering using CD-HIT v4.8.1^75^ ‘cd-hit-est’ with word size 5, sequence identity threshold 0.8 and length difference cut-off 0.8. MGE alignment figures were generated using Easyfig v2.2.3^76^.

### Comparing SDSE and S. pyogenes pangenome

The accessory gene classification scheme described for SDSE and COG annotations were mapped to the *S. pyogenes* pangenome. The merged pangenome was then used to map functional classes of core genes and accessory gene MGE classes across species. Panaroo^69^ combines a small number of genes/coding sequences when merging pangenomes resulting in slightly fewer pangenome genes in the merged pangenome compared to that for the individual species.

Metabolic differences between SDSE and *S. pyogenes* were inferred by searching for well-defined complete KEGG modules present in one species but not the other. Non-MGE and core genes unique to each species were extracted from the pangenomes of each respective species and KO identifiers were assigned using eggNOG-mapper v2.1.7^70^. KO numbers of unique genes from both species were then mapped simultaneously to KEGG modules using KEGG mapper Reconstruct^77^.

### Recombination detection across species

Gene alignments of the 1,116 shared core genes of SDSE and *S. pyogenes* with <25% gap characters were created and analysed with fastGEAR^35^. Interspecies recombination was inferred by two criteria from the fastGEAR results. Lineages or sequence clusters inferred by fastGEAR could contain sequences from one species or both. A predicted recombination event from a cluster containing only one species to a genome from the other species, an event classified as ‘recent’ by fastGEAR, was classified as a putatively cross-species. The presence of both species within the same lineage or cluster when more than one lineage was predicted, could occur in the setting of cross-species recombination of the whole gene or with shared ancestry. As these could not be easily separated, cross-species clusters were recorded separately, and locus tags are provided in Supplementary Table 2b for individual interrogation.

### Vaccine antigen screen

*S. pyogenes* vaccine candidates were screened for presence and sequence diversity in SDSE. Vaccine candidates and screening methods were adapted from a previous report of vaccine antigenic diversity in *S. pyogenes* and updated with new multivalent antigens and formulations^22, 42^ (Supplementary Table 5). The presence of vaccine antigen genes was determined using screen_assembly v1.2.7^22^ with a cut-off of 70% nucleotide identity and 70% coverage. Sequence diversity was presented as nucleotide divergence calculated by BLASTn or predicted amino acid divergence using tBLASTn as indicated.

For the 30-valent M protein vaccine, the 180bp hypervariable 5’ sequence was extracted from publicly available databases and compared against the N-terminal sequence of SDSE *emm* types represented in the 501 genomes from this report at 70% nucleotide identity and 70% coverage. The representative/type SDSE *emm* sequence (e.g., *stG*62647.0, *stG*840.0) was used for the comparison. For the T antigens, nucleotide sequences of the individual subdomains from different T alleles included in the fusion multivalent vaccine formulations were extracted and searched at a threshold of 70% nucleotide identity and 70% coverage. Results were presented as presence/absence as M and T antigens are hypervariable.

The small peptide antigens J8.0, StrepInCor ‘common’ overlapping B and T cell epitope, P*17 and S2 were screened using a six-frame translation of the target genome and search at 100% identity and coverage. As P*17 had two amino acid substitutions at positions 13 and 17 introduced which are not naturally occurring, amino acids at these positions were replaced with a wildcard for the search.

### Data availability

Accessions for raw sequencing data are available in Supplementary Table 1a. The complete genome sequence of *S. dysgalactiae* subsp. *equisimilis* NS3396 was deposited to GenBank (accession CP128987).

### Code availability

Supplementary code used to extract and classify accession segments and MGEs is available at https://github.com/OuliXie/Global_SDSE.

## Author Contributions

OX, DJM and MRD planned the study. RJT, LS, KSS, TH, PG, ACS, MRB, BWB, MDP, MR, DEB, BJC and DJM provided samples and metadata. OX, JMM, AJH, MGJ, JA Lees, NLBZ, OB, SLB, GPC, GTH, LM, JA Lacey, TBJ, SAB, GD, SDB, MJW, SYCT, DJM and MRD designed experimental procedures and generated data. OX, JMM, AJH, MGJ, JA Lees, NLBZ, SLB, GPC, SYCT, DJM and MRD analysed data. OX, JMM, AJH, SLB, GPC, DJM and MRD wrote the manuscript. All authors revised and approved the manuscript.

## Acknowledgements

The work was supported by the National Health and Medical Research Council of Australia (NHMRC) and The Wellcome Trust, UK. MRD was supported by a NHMRC postdoctoral training fellowship (635250) and a University of Melbourne CR Roper Fellowship. OX was supported by the NHMRC postgraduate scholarship (GNT2013831) and Avant Foundation Doctors in Training Research Scholarship (2021/000017). We acknowledge the assistance of the sequencing and pathogen informatics core teams at the Wellcome Sanger Institute, UK where this work was supported by the Wellcome Trust core grants 206194 and 108413/A/15/D.

## References

1. Wright CM, Moorin R, Pearson G et al. Invasive Infections Caused by Lancefield Groups C/G and A *Streptococcus*, Western Australia, Australia, 2000-2018. Emerg Infect Dis 2022; 28: 2190–7.

2. Couture-Cossette A, Carignan A, Mercier A et al. Secular trends in incidence of invasive beta-hemolytic streptococci and efficacy of adjunctive therapy in Quebec, Canada, 1996-2016. PLoS One 2018; 13: e0206289.

3. Sylvetsky N, Raveh D, Schlesinger Y et al. Bacteremia due to beta-hemolytic *Streptococcus* group G: increasing incidence and clinical characteristics of patients. Am J Med 2002; 112: 622–6.

4. Oppegaard O, Mylvaganam H, Kittang BR. Beta-haemolytic group A, C and G streptococcal infections in Western Norway: a 15-year retrospective survey. Clin Microbiol Infect 2015; 21: 171–8.

5. Lambertsen LM, Ingels H, Schønheyder HC et al. Nationwide laboratory-based surveillance of invasive beta-haemolytic streptococci in Denmark from 2005 to 2011. Clin Microbiol Infect 2014; 20: O216–23.

6. Harris P, Siew DA, Proud M et al. Bacteraemia caused by beta-haemolytic streptococci in North Queensland: changing trends over a 14-year period. Clin Microbiol Infect 2011; 17: 1216–22.

7. Shinohara K, Murase K, Tsuchido Y et al. Clonal Expansion of Multidrug-Resistant *Streptococcus dysgalactiae* subspecies *equisimilis* Causing Bacteremia, Japan, 2005-2021. Emerg Infect Dis 2023; 29: 528–39.

8. Broyles LN, Van Beneden C, Beall B et al. Population-based study of invasive disease due to beta-hemolytic streptococci of groups other than A and B. Clin Infect Dis 2009; 48: 706–12.

9. Brandt CM, Spellerberg B. Human infections due to *Streptococcus dysgalactiae* subspecies *equisimilis*. Clin Infect Dis 2009; 49: 766–72.

10. Takahashi T, Sunaoshi K, Sunakawa K et al. Clinical aspects of invasive infections with *Streptococcus dysgalactiae* ssp. *equisimilis* in Japan: differences with respect to *Streptococcus pyogenes* and *Streptococcus agalactiae* infections. Clin Microbiol Infect 2010; 16: 1097–103.

11. McMillan DJ, Bessen DE, Pinho M et al. Population genetics of *Streptococcus dysgalactiae* subspecies *equisimilis* reveals widely dispersed clones and extensive recombination. PLoS One 2010; 5: e11741.

12. McNeilly CL, McMillan DJ. Horizontal gene transfer and recombination in *Streptococcus dysgalactiae* subsp. *equisimilis*. Front Microbiol 2014; 5: 676.

13. Vähäkuopus S, Vuento R, Siljander T et al. Distribution of *emm* types in invasive and non-invasive group A and G streptococci. Eur J Clin Microbiol Infect Dis 2012; 31: 1251–6.

14. Pinho MD, Melo-Cristino J, Ramirez M. Fluoroquinolone resistance in *Streptococcus dysgalactiae* subsp. *equisimilis* and evidence for a shared global gene pool with *Streptococcus pyogenes*. Antimicrob Agents Chemother 2010; 54: 1769–77.

15. van Sorge NM, Cole JN, Kuipers K et al. The classical lancefield antigen of group A *Streptococcus* is a virulence determinant with implications for vaccine design. Cell Host Microbe 2014; 15: 729–40.

16. Beres SB, Zhu L, Pruitt L et al. Integrative Reverse Genetic Analysis Identifies Polymorphisms Contributing to Decreased Antimicrobial Agent Susceptibility in *Streptococcus pyogenes*. mBio 2022; 13: e0361821.

17. Ahmad Y, Gertz RE, Jr., Li Z et al. Genetic relationships deduced from emm and multilocus sequence typing of invasive *Streptococcus dysgalactiae* subsp. *equisimilis* and *S. canis* recovered from isolates collected in the United States. J Clin Microbiol 2009; 47: 2046–54.

18. Chochua S, Rivers J, Mathis S et al. Emergent Invasive Group A *Streptococcus dysgalactiae* subsp. *equisimilis*, United States, 2015-2018. Emerg Infect Dis 2019; 25: 1543–7.

19. Chochua S, Metcalf BJ, Li Z et al. Population and Whole Genome Sequence Based Characterization of Invasive Group A Streptococci Recovered in the United States during 2015. mBio 2017; 8: e01422–17.

20. Davies MR, Shera J, Van Domselaar GH et al. A novel integrative conjugative element mediates genetic transfer from group G *Streptococcus* to other beta-hemolytic Streptococci. J Bacteriol 2009; 191: 2257–65.

21. Palmieri C, Magi G, Creti R et al. Interspecies mobilization of an *ermT*-carrying plasmid of *Streptococcus dysgalactiae* subsp. *equisimilis* by a coresident ICE of the ICESa2603 family. J Antimicrob Chemother 2013; 68: 23–6.

22. Davies MR, McIntyre L, Mutreja A et al. Atlas of group A streptococcal vaccine candidates compiled using large-scale comparative genomics. Nature Genetics 2019; 51: 1035–43.

23. Lother SA, Demczuk W, Martin I et al. Clonal Clusters and Virulence Factors of Group C and G *Streptococcus* Causing Severe Infections, Manitoba, Canada, 2012-2014. Emerg Infect Dis 2017; 23: 1079–88.

24. Oppegaard O, Mylvaganam H, Skrede S et al. Emergence of a *Streptococcus dysgalactiae* subspecies *equisimilis* stG62647-lineage associated with severe clinical manifestations. Sci Rep 2017; 7: 7589.

25. Kaci A, Jonassen CM, Skrede S et al. Genomic epidemiology of *Streptococcus dysgalactiae* subsp. *equisimilis* strains causing invasive disease in Norway during 2018. Frontiers in Microbiology 2023; 14.

26. Dale JB, Walker MJ. Update on group A streptococcal vaccine development. Curr Opin Infect Dis 2020; 33: 244–50.

27. Vekemans J, Gouvea-Reis F, Kim JH et al. The Path to Group A *Streptococcus* Vaccines: World Health Organization Research and Development Technology Roadmap and Preferred Product Characteristics. Clin Infect Dis 2019; 69: 877–83.

28. Davies MR, McMillan DJ, Van Domselaar GH et al. Phage 3396 from a *Streptococcus dysgalactiae* subsp. *equisimilis* pathovar may have its origins in *Streptococcus pyogenes*. J Bacteriol 2007; 189: 2646–52.

29. Lees JA, Harris SR, Tonkin-Hill G et al. Fast and flexible bacterial genomic epidemiology with PopPUNK. Genome Res 2019; 29: 304–16.

30. Didelot X, Croucher NJ, Bentley SD et al. Bayesian inference of ancestral dates on bacterial phylogenetic trees. Nucleic Acids Res 2018; 46: e134.

31. Croucher NJ, Page AJ, Connor TR et al. Rapid phylogenetic analysis of large samples of recombinant bacterial whole genome sequences using Gubbins. Nucleic Acids Res 2015; 43: e15.

32. Ikebe T, Okuno R, Sasaki M et al. Molecular characterization and antibiotic resistance of *Streptococcus dysgalactiae* subspecies *equisimilis* isolated from patients with streptococcal toxic shock syndrome. J Infect Chemother 2018; 24: 117–22.

33. Beres SB, Olsen RJ, Long SW et al. Analysis of the Genomics and Mouse Virulence of an Emergent Clone of *Streptococcus dysgalactiae* subspecies *equisimilis*. Microbiol Spectr 2023: e0455022.

34. Oppegaard O, Mylvaganam H, Skrede S et al. Exploring the arthritogenicity of *Streptococcus dysgalactiae* subspecies *equisimilis*. BMC Microbiology 2018; 18: 17.

35. Mostowy R, Croucher NJ, Andam CP et al. Efficient Inference of Recent and Ancestral Recombination within Bacterial Populations. Mol Biol Evol 2017; 34: 1167–82.

36. Khedkar S, Smyshlyaev G, Letunic I et al. Landscape of mobile genetic elements and their antibiotic resistance cargo in prokaryotic genomes. Nucleic Acids Research 2022; 50: 3155–68.

37. McShan WM, McCullor KA, Nguyen SV. The Bacteriophages of *Streptococcus pyogenes*. Microbiol Spectr 2019; 7.

38. Berbel D, Càmara J, González-Díaz A et al. Deciphering mobile genetic elements disseminating macrolide resistance in *Streptococcus pyogenes* over a 21 year period in Barcelona, Spain. J Antimicrob Chemother 2021; 76: 1991–2003.

39. Arredondo-Alonso S, Gladstone RA, Pöntinen AK et al. Mge-cluster: a reference-free approach for typing bacterial plasmids. NAR Genom Bioinform 2023; 5: lqad066.

40. Sumby P, Barbian KD, Gardner DJ et al. Extracellular deoxyribonuclease made by group A *Streptococcus* assists pathogenesis by enhancing evasion of the innate immune response. Proc Natl Acad Sci U S A 2005; 102: 1679–84.

41. Sumby P, Porcella SF, Madrigal AG et al. Evolutionary origin and emergence of a highly successful clone of serotype M1 group A *Streptococcus* involved multiple horizontal gene transfer events. J Infect Dis 2005; 192: 771–82.

42. Harbison-Price N, Rivera-Hernandez T, Osowicki J et al. Current Approaches to Vaccine Development of *Streptococcus pyogenes*. In: Ferretti JJ, Stevens DL, Fischetti VA, eds. Streptococcus pyogenes: Basic Biology to Clinical Manifestations. Oklahoma City (OK): University of Oklahoma Health Sciences Center © The University of Oklahoma Health Sciences Center., 2022.

43. Porcellato D, Smistad M, Skeie SB et al. Whole genome sequencing reveals possible host species adaptation of *Streptococcus dysgalactiae*. Sci Rep 2021; 11: 17350.

44. Diop A, Torrance EL, Stott CM et al. Gene flow and introgression are pervasive forces shaping the evolution of bacterial species. Genome Biol 2022; 23: 239.

45. Davies MR, Keller N, Brouwer S et al. Detection of *Streptococcus pyogenes* M1(UK) in Australia and characterization of the mutation driving enhanced expression of superantigen SpeA. Nat Commun 2023; 14: 1051.

46. Nordström T, Malcolm J, Magor G et al. In vivo efficacy of a chimeric peptide derived from the conserved region of the M protein against group C and G streptococci. Clin Vaccine Immunol 2012; 19: 1984–7.

47. Ishihara H, Ogura K, Miyoshi-Akiyama T et al. Prevalence and genomic characterization of Group A *Streptococcus dysgalactiae* subsp. *equisimilis* isolated from patients with invasive infections in Toyama prefecture, Japan. Microbiol Immunol 2020; 64: 113–22.

48. Shimomura Y, Okumura K, Murayama SY et al. Complete genome sequencing and analysis of a Lancefield group G *Streptococcus dysgalactiae* subsp. *equisimilis* strain causing streptococcal toxic shock syndrome (STSS). BMC Genomics 2011; 12: 17.

49. Okumura K, Shimomura Y, Murayama SY et al. Evolutionary paths of streptococcal and staphylococcal superantigens. BMC Genomics 2012; 13: 404.

50. Brandt CM, Haase G, Schnitzler N et al. Characterization of blood culture isolates of *Streptococcus dysgalactiae* subsp. *equisimilis* possessing Lancefield’s group A antigen. J Clin Microbiol 1999; 37: 4194–7.

51. Babbar A, Nitsche-Schmitz DP, Pieper DH et al. Draft Genome Sequence of *Streptococcus dysgalactiae* subsp. *equisimilis* Strain C161L1 Isolated in Vellore, India. Genome Announc 2017; 5.

52. Wang X, Zhang X, Zong Z. Genome sequence and virulence factors of a group G *Streptococcus dysgalactiae* subsp. *equisimilis* strain with a new element carrying *erm(B)*. Sci Rep 2016; 6: 20389.

53. Rebelo AR, Bortolaia V, Leekitcharoenphon P et al. One Day in Denmark: Comparison of Phenotypic and Genotypic Antimicrobial Susceptibility Testing in Bacterial Isolates From Clinical Settings. Front Microbiol 2022; 13: 804627.

54. Cuénod A, Foucault F, Pflüger V et al. Factors Associated With MALDI-TOF Mass Spectral Quality of Species Identification in Clinical Routine Diagnostics. Front Cell Infect Microbiol 2021; 11: 646648.

55. Jagne I, Keeley AJ, Bojang A et al. Impact of intra-partum azithromycin on carriage of group A *Streptococcus* in the Gambia: a posthoc analysis of a double-blind randomized placebo-controlled trial. BMC Infect Dis 2022; 22: 103.

56. Wood DE, Lu J, Langmead B. Improved metagenomic analysis with Kraken 2. Genome Biol 2019; 20: 257.

57. Prjibelski A, Antipov D, Meleshko D et al. Using SPAdes De Novo Assembler. Curr Protoc Bioinformatics 2020; 70: e102.

58. Seemann T. Prokka: rapid prokaryotic genome annotation. Bioinformatics 2014; 30: 2068–9.

59. Jolley KA, Bray JE, Maiden MCJ. Open-access bacterial population genomics: BIGSdb software, the PubMLST.org website and their applications. Wellcome Open Res 2018; 3: 124.

60. Baines SL, Howden BP, Heffernan H et al. Rapid Emergence and Evolution of Staphylococcus aureus Clones Harboring fusC-Containing Staphylococcal Cassette Chromosome Elements. Antimicrob Agents Chemother 2016; 60: 2359–65.

61. Page AJ, Ainsworth EV, Langridge GC. socru: typing of genome-level order and orientation around ribosomal operons in bacteria. Microb Genom 2020; 6.

62. Lefort V, Desper R, Gascuel O. FastME 2.0: A Comprehensive, Accurate, and Fast Distance-Based Phylogeny Inference Program. Mol Biol Evol 2015; 32: 2798–800.

63. Minh BQ, Schmidt HA, Chernomor O et al. IQ-TREE 2: New Models and Efficient Methods for Phylogenetic Inference in the Genomic Era. Mol Biol Evol 2020; 37: 1530–4.

64. Smith MR. Information theoretic generalized Robinson-Foulds metrics for comparing phylogenetic trees. Bioinformatics 2020; 36: 5007–13.

65. Nascimento M, Sousa A, Ramirez M et al. PHYLOViZ 2.0: providing scalable data integration and visualization for multiple phylogenetic inference methods. Bioinformatics 2017; 33: 128–9.

66. Liu B, Zheng D, Zhou S, et al. VFDB 2022: a general classification scheme for bacterial virulence factors. Nucleic Acids Res 2022; 50: D912–d7.

67. Jespersen MG, Hayes A, Davies MR. Magphi: Sequence extraction tool from FASTA and GFF3 files using seed pairs. Journal of Open Source Software 2022; 7: 4369.

68. Feldgarden M, Brover V, Gonzalez-Escalona N et al. AMRFinderPlus and the Reference Gene Catalog facilitate examination of the genomic links among antimicrobial resistance, stress response, and virulence. Sci Rep 2021; 11: 12728.

69. Tonkin-Hill G, MacAlasdair N, Ruis C et al. Producing polished prokaryotic pangenomes with the Panaroo pipeline. Genome Biol 2020; 21: 180.

70. Cantalapiedra CP, Hernández-Plaza A, Letunic I et al. eggNOG-mapper v2: Functional Annotation, Orthology Assignments, and Domain Prediction at the Metagenomic Scale. Mol Biol Evol 2021; 38: 5825–9.

71. Jespersen MG, Hayes A, Davies MR. Corekaburra: pan-genome post-processing using core gene synteny. Journal of Open Source Software 2022; 7: 4910.

72. Mistry J, Finn RD, Eddy SR et al. Challenges in homology search: HMMER3 and convergent evolution of coiled-coil regions. Nucleic Acids Res 2013; 41: e121.

73. Abby SS, Cury J, Guglielmini J et al. Identification of protein secretion systems in bacterial genomes. Sci Rep 2016; 6: 23080.

74. Katoh K, Standley DM. MAFFT multiple sequence alignment software version 7: improvements in performance and usability. Mol Biol Evol 2013; 30: 772–80.

75. Fu L, Niu B, Zhu Z et al. CD-HIT: accelerated for clustering the next-generation sequencing data. Bioinformatics 2012; 28: 3150–2.

76. Sullivan MJ, Petty NK, Beatson SA. Easyfig: a genome comparison visualizer. Bioinformatics 2011; 27: 1009–10.

77. Kanehisa M, Sato Y, Kawashima M. KEGG mapping tools for uncovering hidden features in biological data. Protein Sci 2022; 31: 47–53.

