## Supplementary Materials for "Inter-species gene flow drives ongoing evolution of *Streptococcus pyogenes* and *Streptococcus dysgalactiae* subsp. *equisimilis*"

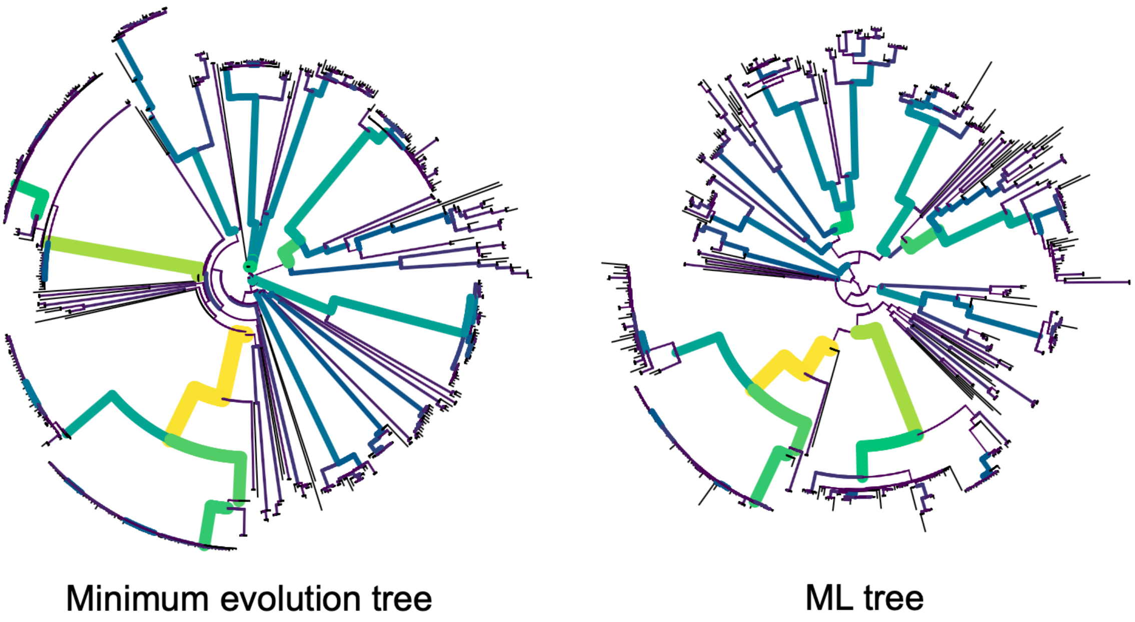


**Supplementary Figure 1.** Topological comparison of a *Streptococcus dysgalactiae* subsp. *equisimilis* minimum evolution phylogeny using recombination masked genomic distances with a maximum likelihood (ML) phylogeny constructed without correction for recombination. ML tree was derived from 102,866 core SNPS and 74,069 parsimony-informative sites aligned to the GGS_124 reference genome (accession NC_012891.1) and inferred using a GTR + G4 model. Matched, shared branches are bolded and coloured. The topologies demonstrated a similar internal branching structure radiating into comparable major clusters but with rearrangements in the deepest ancestral branches. Tips were shortened when recombination was masked consistent with the effect of recombination on branch lengths. The information-based generalised Robinson-Foulds distance using mutual clustering information, where 1 represents the maximum distance between trees, was 0.28 and consistent with general agreement of the tree topology between the phylogenies.

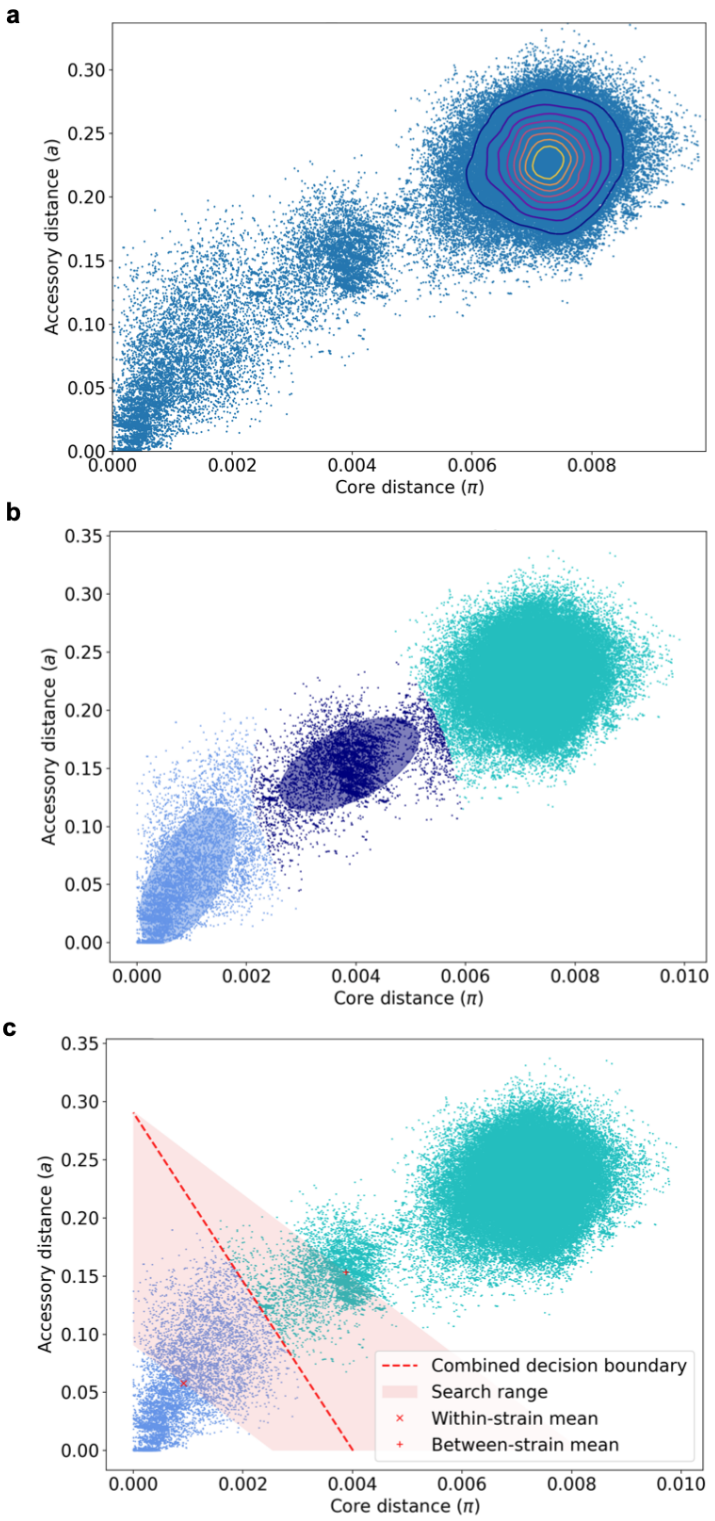


**Supplementary Figure 2.** PopPUNK model fitting of the global 501 *Streptococcus dysgalactiae* subsp. *equisimilis* genomes. **a)** Scatterplot of core and accessory distances for all pair-wise genome comparisons. **b)** Fit of 3 spatial clusters to pair-wise distances using

Bayesian Gaussian Mixture Model. **c)** 2D model boundary refinement to determine within cluster and between cluster distance boundaries.

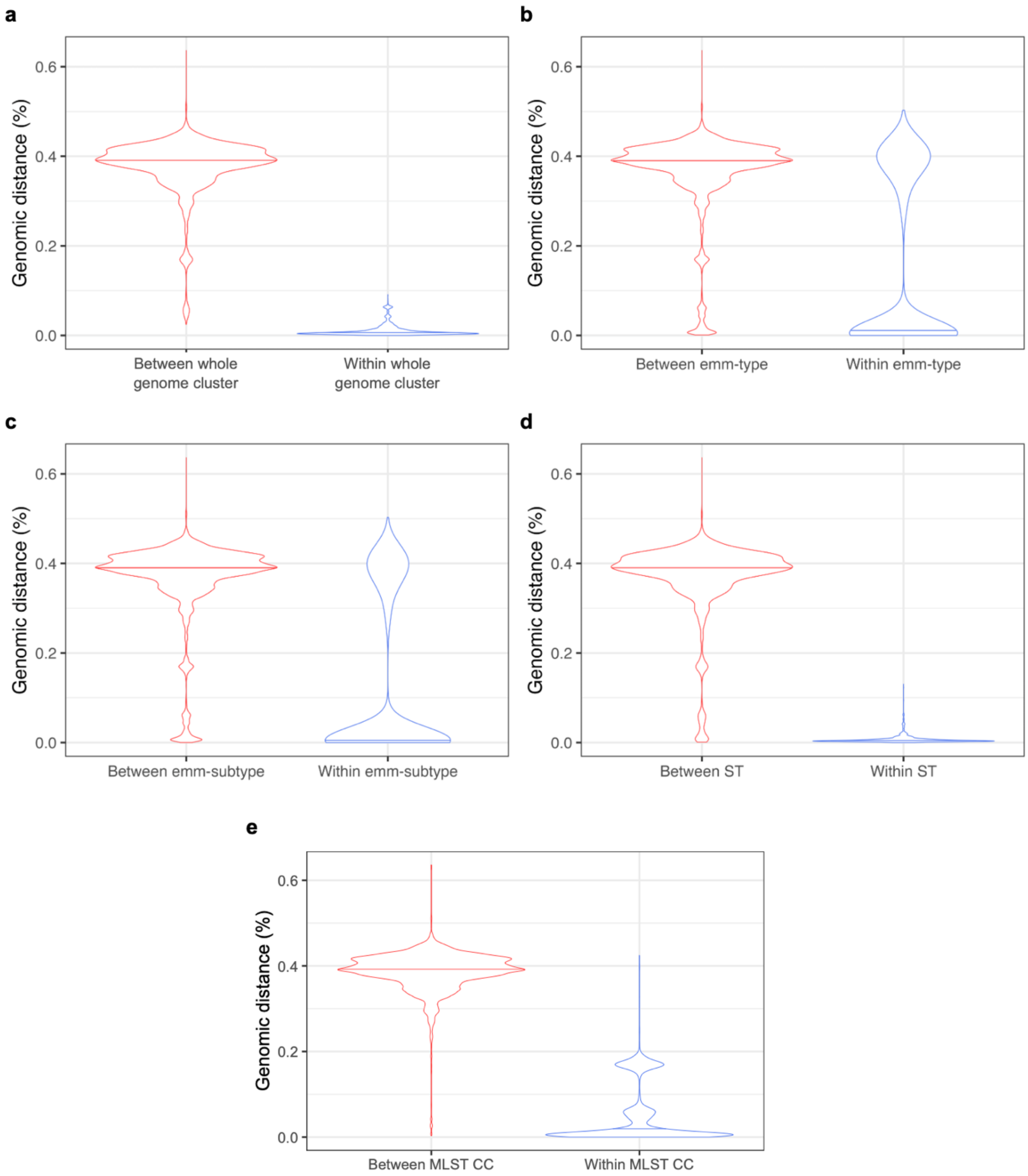


**Supplementary Figure 3.** Violin plots of within vs between cluster recombination-masked genomic distances (100% – average nucleotide identity). Median indicated by horizontal lines within violin plots. **a)** Whole genome clusters defined by PopPUNK. **b)** *emm* type. **c)** *emm* subtype. **d)** MLST. **e)** Multilocus sequence type (MLST) single locus variant clonal complexes (CC). *emm* and *emm* subtype distances demonstrated a bimodal distribution, indicating presence of the same *emm* or *emm* subtype across different genomic backgrounds. MLST was better able to distinguish different genomic backgrounds but split otherwise very similar isolates. MLST CC identified some closely related isolates which were split by MLST, but CCs were also present across distinct genetic backgrounds.

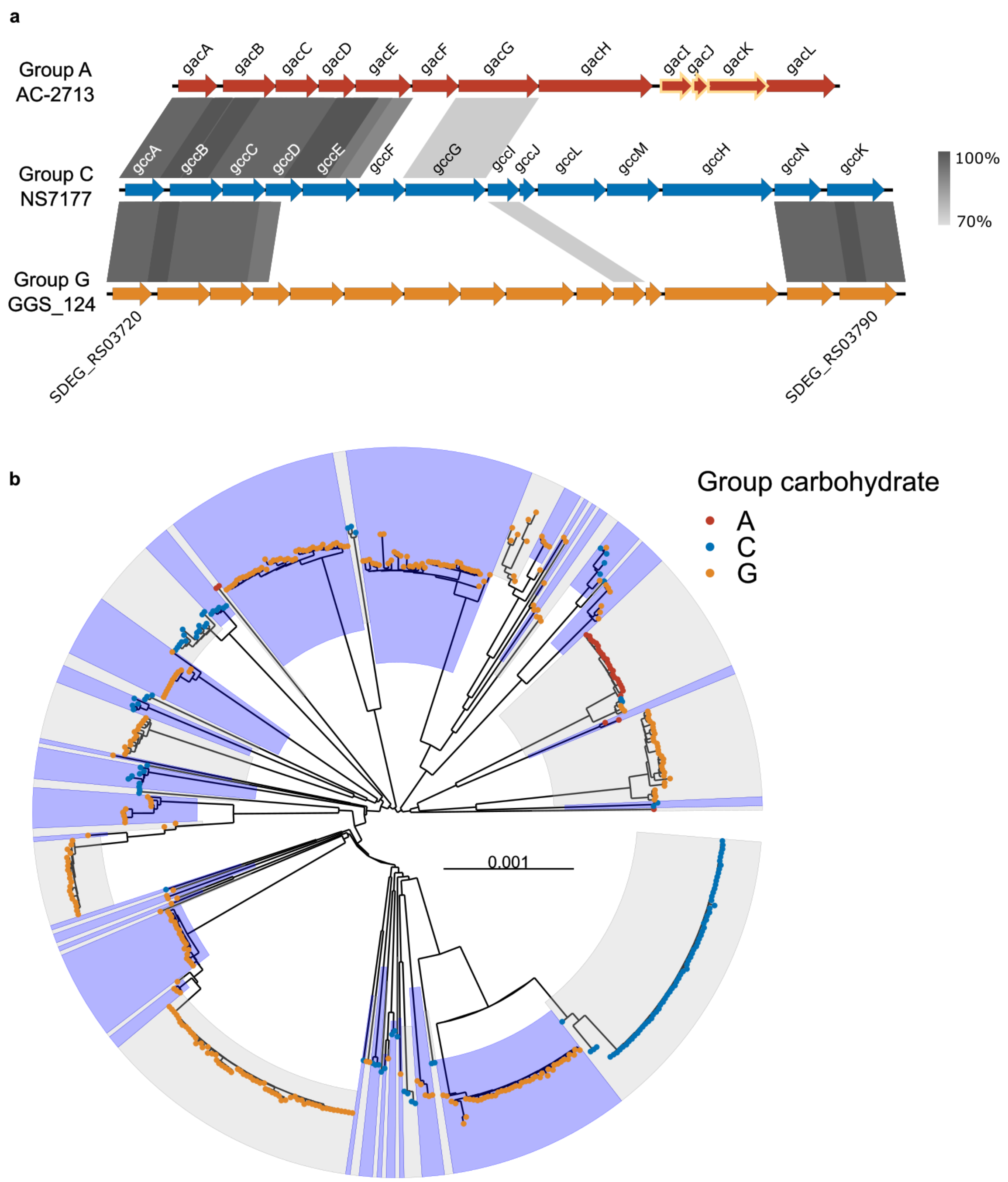


**Supplementary Figure 4.** The Lancefield carbohydrate loci of group A, C and G *Streptococcus dysgalactiae* subsp. *equisimilis*. **a)** Pairwise alignment of Lancefield group carbohydrate synthesis loci from representative group A, C, and G *Streptococcus dysgalactiae* subsp. *equisimilis* (SDSE) genomes. Gene architecture and comparison of group A (AC-2713, accession NC_019042.1), group C (NS7177, accession ERR026931) and group G (GGS_124, accession NC_012891.1) SDSE carbohydrate synthesis loci. Regions of genomic similarity were inferred using tBLASTx and plotted using Easyfig v2.2.3[76]. The grey gradient in the legend refers to the Blast identity percentage. Gene names are labelled as described previously for the group A[15] and group C[18] carbohydrate locus. Genes names for the group G carbohydrate locus are labelled with locus tags from reference genome GGS_124. The genes essential for the immunodominant *N*-acetylglucosamine sidechain for the group A carbohydrate[15], *gacI-gacK,* are outlined in yellow. Genes homologous to *gacA, gacB,* and *gacC* were present across all carbohydrate loci. Putative genes *gccI, gccN* and *gccK* were shared between group C and G loci and *gacD, gacE,* and *gacG* were shared between group A and C loci. The group carbohydrate was predicted based on presence of all genes of a carbohydrate locus in the SDSE pangenome. **b)** Distribution of Lancefield antigen/group carbohydrate across SDSE whole genome clusters. Minimum evolution phylogeny using recombination-masked genomic distances with inferred group A, C, and G carbohydrate indicated by points on the tree tips. Distinct genome clusters are indicated by alternating blue and grey shades from internal nodes. The group G carbohydrate was found in all isolates from 16 genome clusters and group C from 9 genome clusters. The group A carbohydrate, normally associated with *S. pyogenes*, was found in 4 distinct genome clusters, including one cluster with both group C and G isolates.

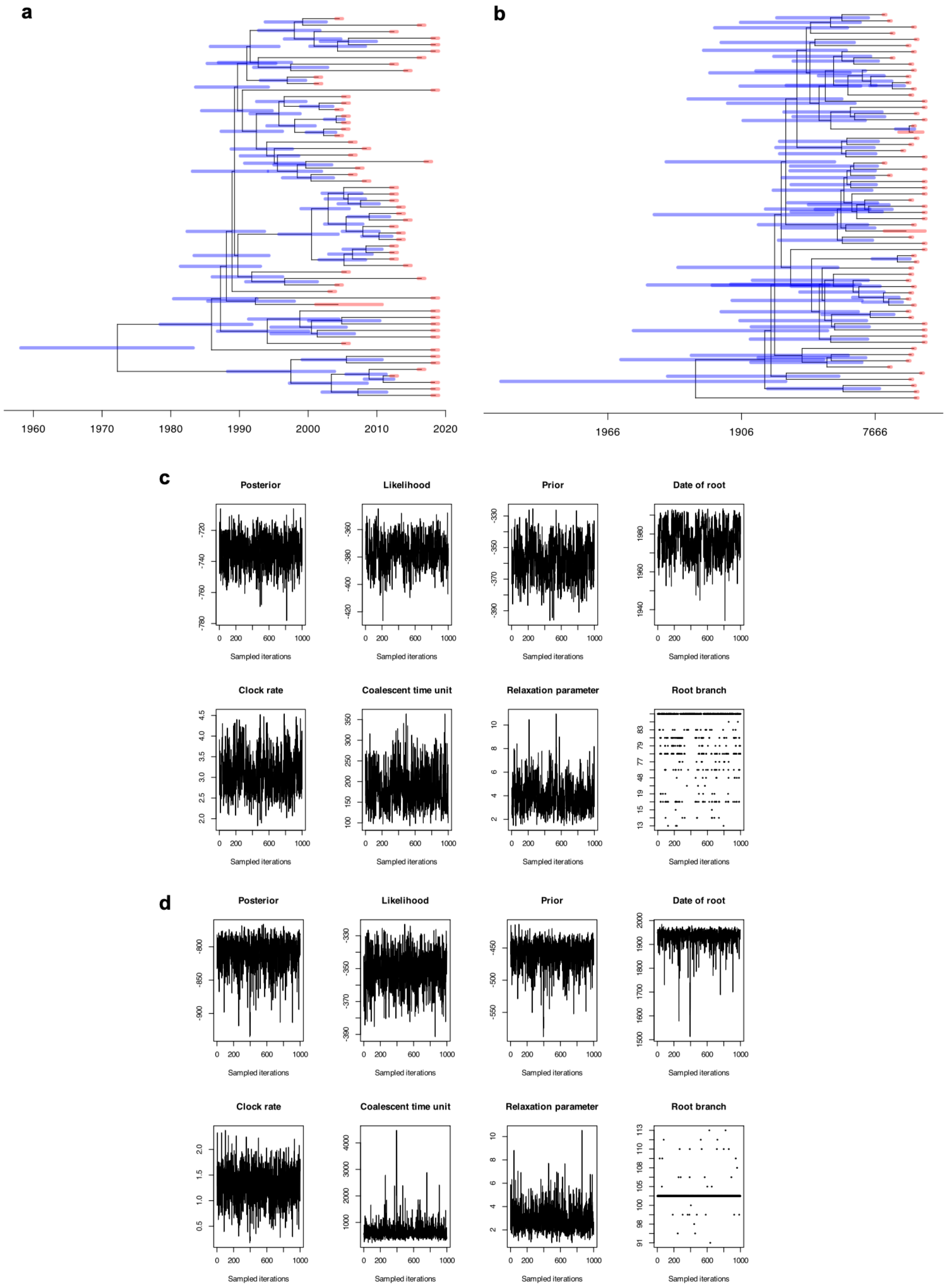

a
1960
1970
1980
1990
2000
2010
2020
b
1966
1906
7666
c
Posterior
Likelihood
Prior
Date of root
Clock rate
Coalescent time unit
Relaxation parameter
Root branch
Sampled iterations
d
Posterior
Likelihood
Prior
Date of root
Clock rate
Coalescent time unit
Relaxation parameter
Root branch
Sampled iterations

**Supplementary Figure 5. a)** Bayesian inference of the most recent common ancestor of 'genome cluster 2', a cluster with isolates found across 7 countries and including *stG*6792 ST 17 isolates which have been described as the most frequent cause of invasive disease in Japan. Dated phylogeny with credible intervals of the nodes indicated by blue bars and dating intervals of the isolates indicated by red bars. The root date was inferred as 1977 (credibility interval 1960 – 1991) suggesting recent emergence of a globally dispersed and disease-causing lineage. **b)** Bayesian inference of the most recent common ancestor of 'genome cluster 1', a cluster with isolates found across 5 countries and consisting of *stG*62647 isolates which have been described as increasing in frequency in Norway. Four non-*stG*62647 isolates were excluded as they were >1500 SNPs distant. The root date was inferred as 1933 (credibility interval 1855 – 1969). **c)** Markov Chain Monte Carlo trace plot for inferred parameters for 'genome cluster 2' and **d)** for 'genome cluster 1'.

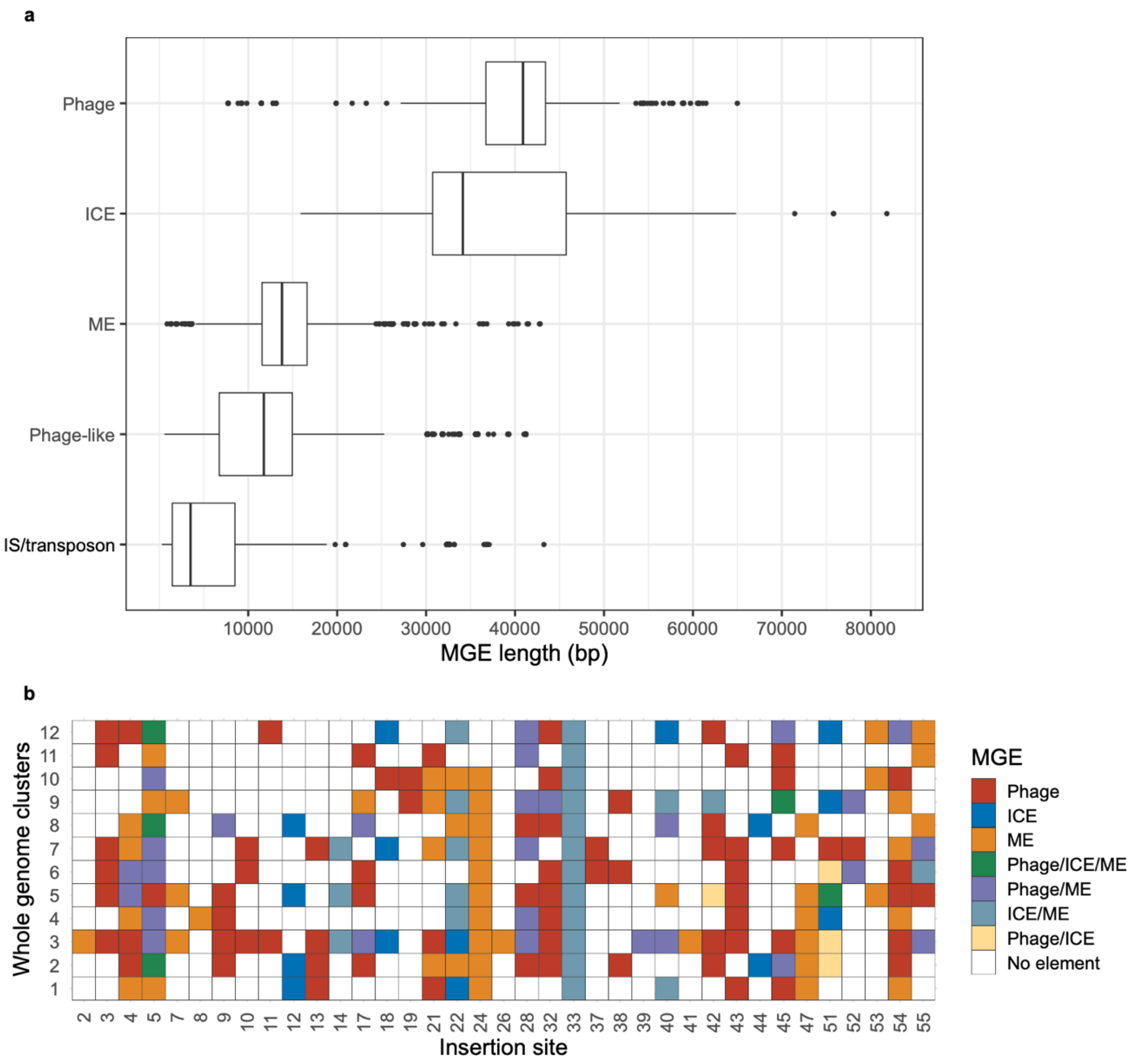


**Supplementary Figure 6. a)** Box and whisker plots of the length of *Streptococcus dysgalactiae* subsp. *equisimilis* (SDSE) mobile genetic element (MGE) classes classified using integrase/recombinase classes and phage/integrative conjugative element (ICE) structural genes on segments of accessory genes. The method was adapted from Khedkar *et al.*[36] and enhanced by the classification and localisation of accessory segments using Corekaburra[71]. Nested elements indicated by the presence of four or more integrase/recombinase genes or co-localisation of multiple phage and ICE genes on the segment were excluded. The range of MGE lengths fell within expected ranges for each class of element. The median length of elements classified as mobility elements (ME) was smaller than phage or ICE and consistent with degraded elements or fragmented phage or ICE. ICE, integrative conjugative element; IS, insertion sequence; ME, mobility element. **b)** Distribution of different MGEs at 37 SDSE chromosomal insertion sites within 12 most sampled whole genome clusters. Only chromosomal insertion sites where at least 5 MGE were observed in the dataset are represented. Prophage and phage-like elements were grouped and IS/transposons were excluded from the analysis. The insertion regions were occupied across phylogroups without any clear patterns of restriction, providing a basis for movement of MGEs across genome clusters.

a

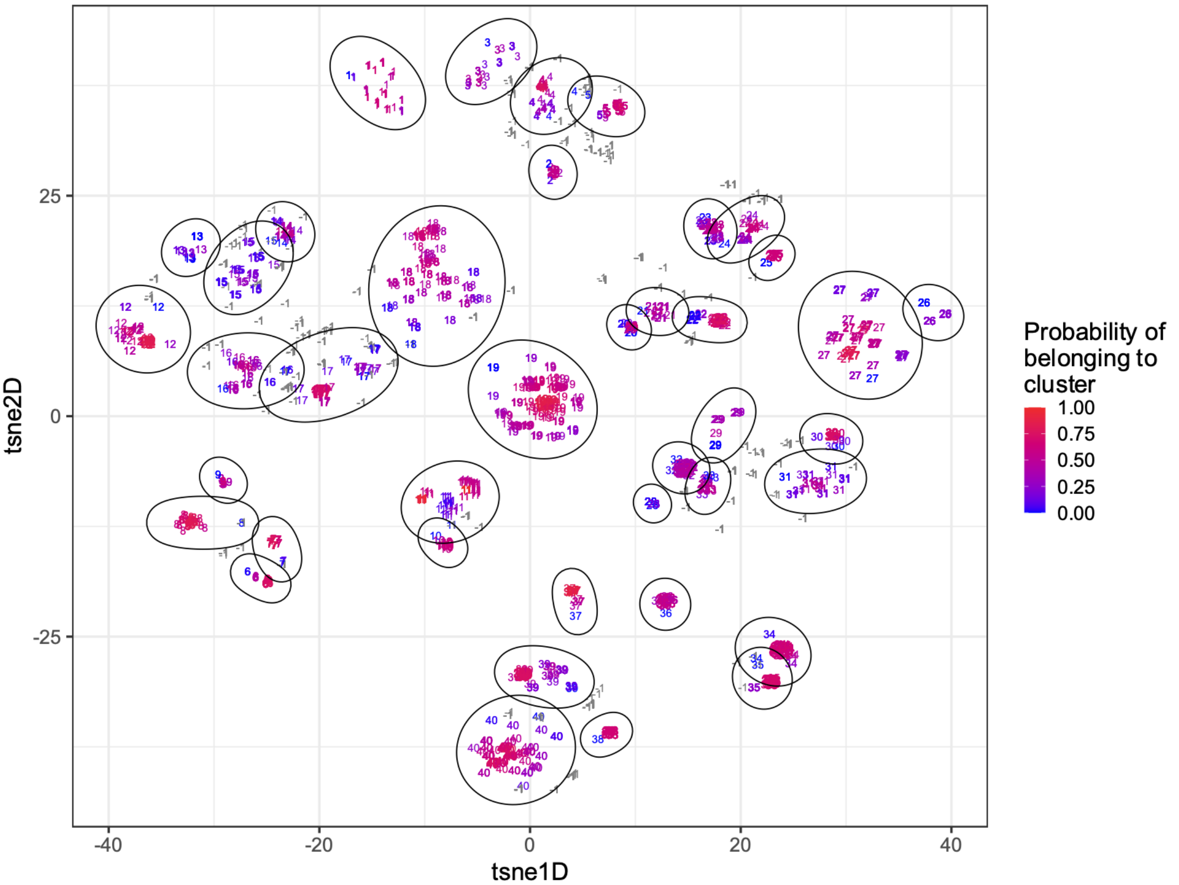

tsne2D
tsne1D
25
0
-25
-40
-20
0
20
40
Probability of belonging to cluster
1.00
0.75
0.50
0.25
0.00


b

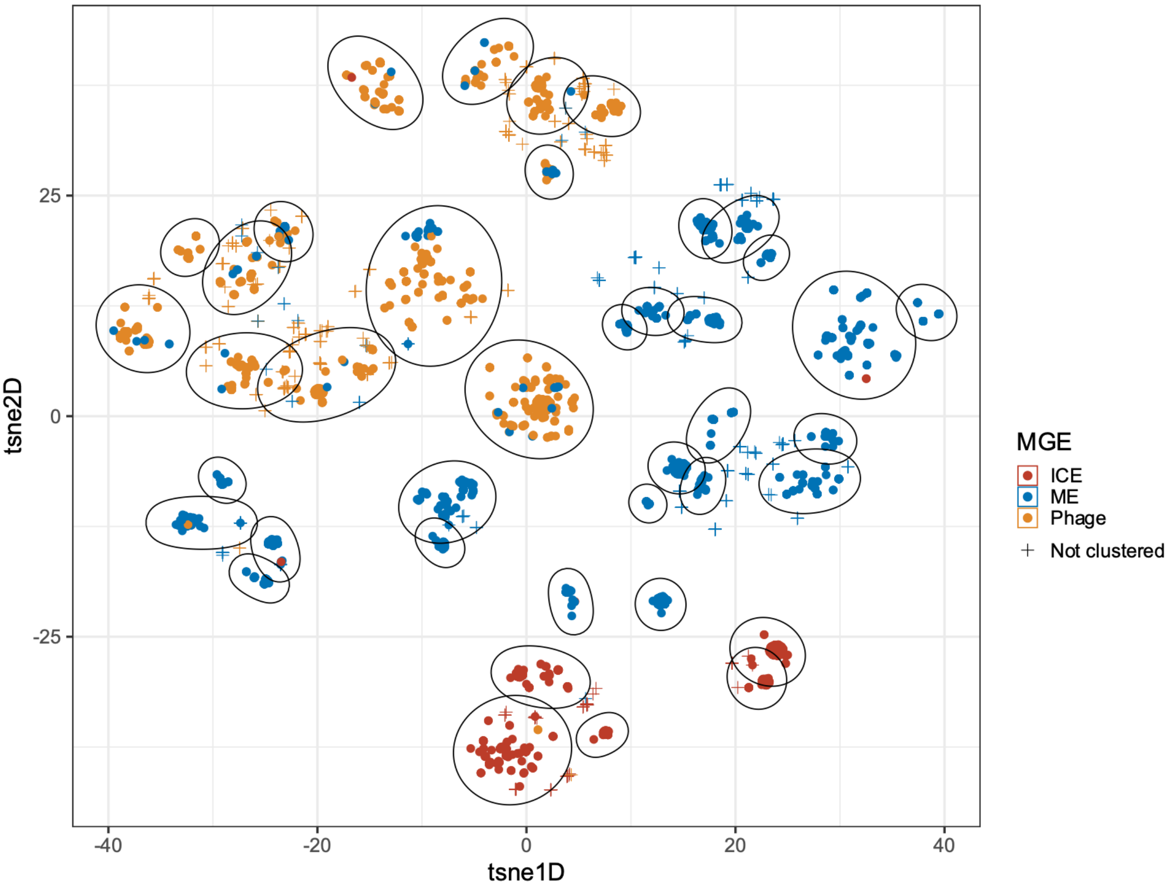

tsne2D
tsne1D
25
0
-25
-40
-20
0
20
40
MGE
ICE
ME
Phage
Not clustered

**Supplementary Figure 7.** Mobile genetic element (MGE) clusters from 16 conserved insertion regions across *Streptococcus dysgalactiae* subsp. *equisimilis* and *S. pyogenes* obtained using mge-cluster[39]. **a)** The probability of each MGE extracted from insertion regions belonging to a cluster. Clusters are denoted by ellipsoids. Segments are labelled by their cluster number with '-1' designated for isolates which were not assigned a cluster. **b)** Clusters coloured by MGE type. Prophage and phage-like elements were grouped together for the analysis. mge-cluster demonstrated excellent separation of element types. As mobility elements (ME) could represent degraded, nested or fragmented elements, clustering of prophage or integrative conjugative elements (ICE) with ME is expected. Two elements annotated as phage in cluster 40 were grouped with ICE and one ICE was clustered with phage elements in cluster 1. Inspection of those elements revealed they were rare MGEs inserted into likely degraded ICE or prophages respectively, at the same insertion region as other elements in that cluster. Therefore, mis-clustering was infrequent and likely caused by presence of residual genes from a degraded MGE. ICE, integrative conjugative element; IS, insertion sequence; ME, mobility element.

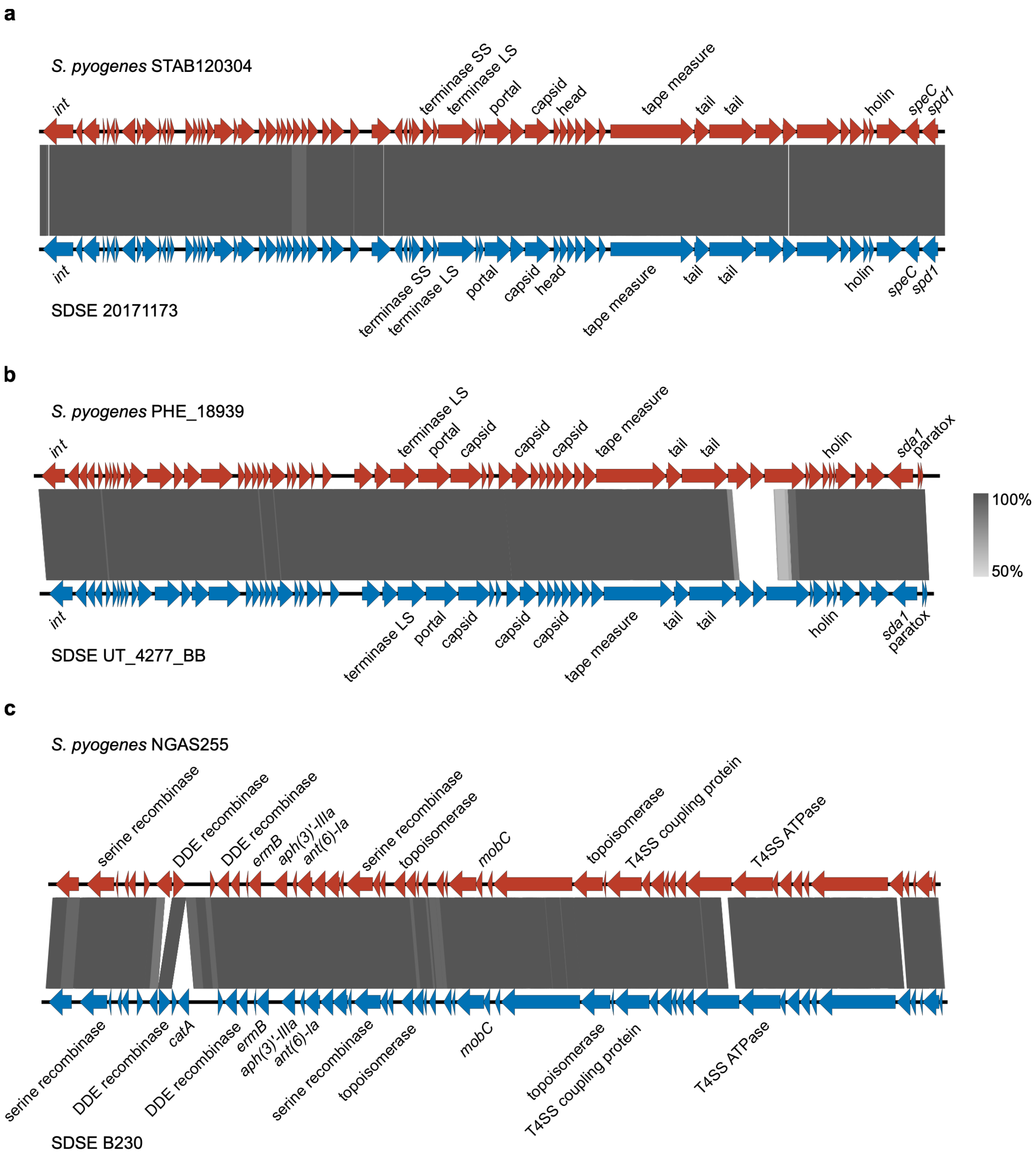


**Supplementary Figure 8.** Pairwise alignment of homologous mobile genetic elements (MGEs) found at shared chromosomal insertion regions between *Streptococcus dysgalactiae* subsp. *equisimilis* (SDSE) and *S. pyogenes*. **a)** Prophage at insertion site 17 carrying the exotoxin gene *speC* and streptodornase gene *spd1*. **b)** Prophage at insertion region 19 carrying an allele of *sda1* with 95% identity to *sda1* found on the M1T1 prophage ϕ5005.3. **c)** Nested integrative conjugative element-like and insertion sequence/transposon element at insertion region 49 carrying multiple AMR genes including the macrolide resistance *ermB* gene, and aminoglycoside resistance genes *ant(6)-Ia* and *aph(3')-IIIa*. The SDSE element has an additional insertion of an insertion sequence/transposon with the chloramphenicol resistance gene *catA*.

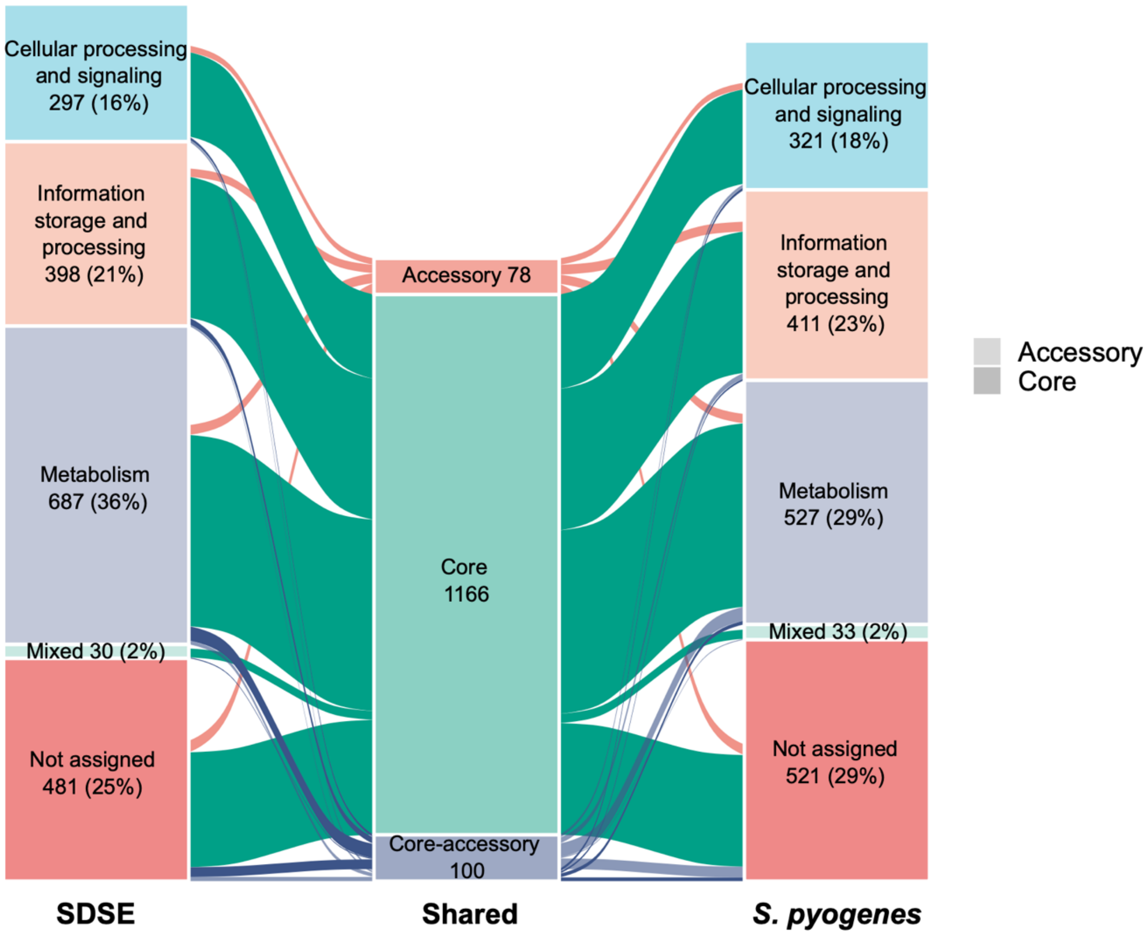


**Supplementary Figure 9.** Alluvial plot of the overlapping core and non-mobile genetic element (MGE) accessory genes between *Streptococcus dysgalactiae* subsp. *equisimilis* (SDSE) and *S. pyogenes* by functional categories. The shared genome was classified into core, core-accessory where a gene was core in one species but accessory in the other, and accessory when it was accessory in both. There was extensive overlap in genes in all functional categories and most were core in both species. More than 50% of genes from each functional category were shared including 68.9% of SDSE and 86.1% of *S. pyogenes* metabolic genes.

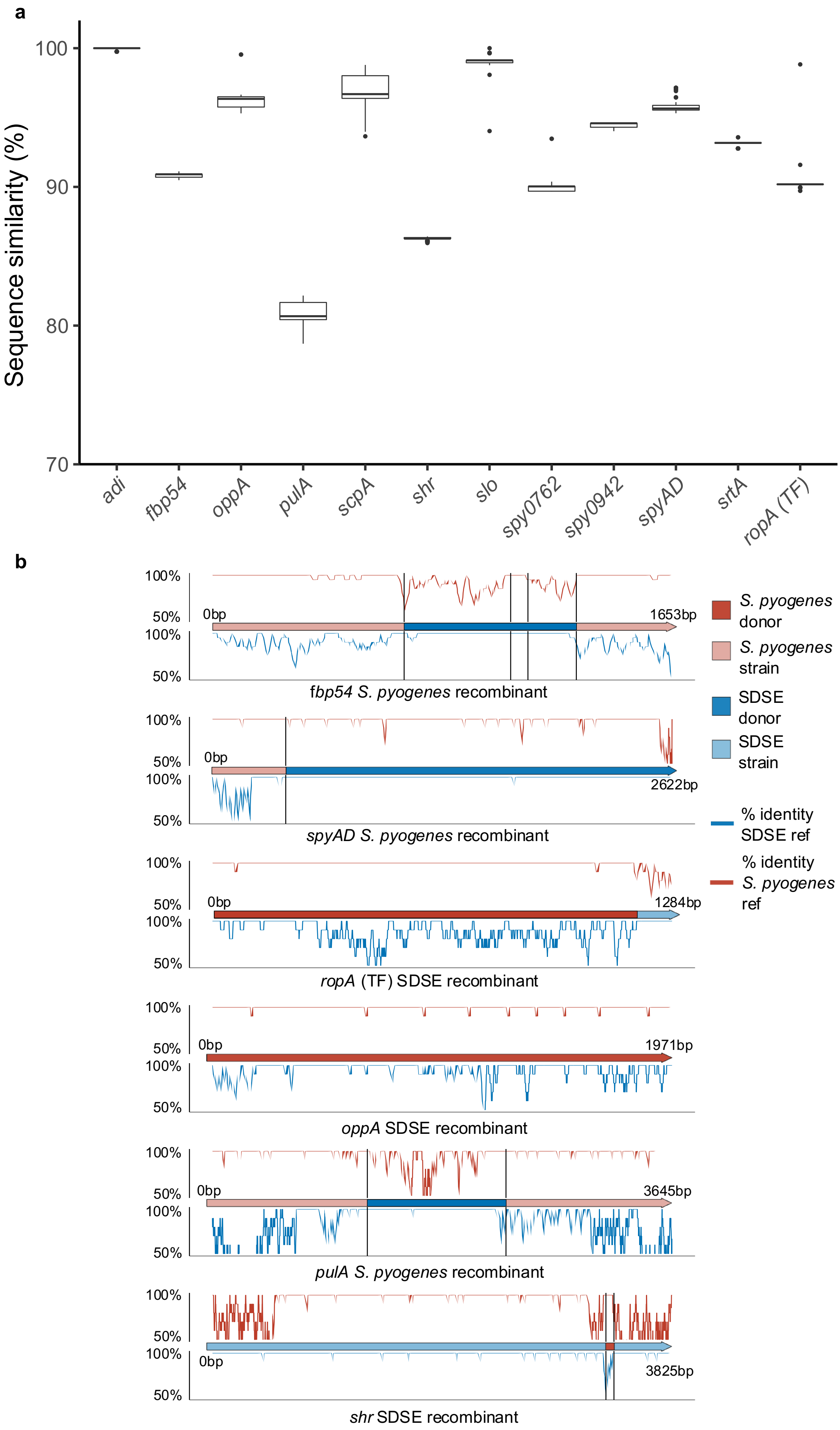

a
Sequence similarity (%)
100
90
80
70
adi
fbp54
oppA
pulA
scpA
shr
slo
spy0762
spy0942
spyAD
srtA
ropA (TF)
b
100%
50%
0bp
1653bp
fbp54 S. pyogenes recombinant
2622bp
spyAD S. pyogenes recombinant
1284bp
ropA (TF) SDSE recombinant
1971bp
oppA SDSE recombinant
3645bp
pulA S. pyogenes recombinant
3825bp
shr SDSE recombinant
S. pyogenes donor
S. pyogenes strain
SDSE donor
SDSE strain
% identity SDSE ref
% identity S. pyogenes ref

**Supplementary Figure 10.** Homology and recombination of shared vaccine candidates across *Streptococcus dysgalactiae* subsp. *equisimilis* and *S. pyogenes*. **a)** Box and whisker plot of amino acid sequence variation of 12 candidate *S. pyogenes* vaccine antigens which were present in >99% of isolates in the global *Streptococcus dysgalactiae* subsp. *equisimilis* (SDSE) (n=501) and *S. pyogenes* (n=2083) population genomic databases. Amino acid similarity was determined using tBLASTn against the *S. pyogenes*-derived reference amino acid sequence[22, 42]. The upper, middle and lower bars of the boxplot indicate the 75% quantile, median and 25% quantile respectively. Outliers are indicated by points greater than the 75% quantile plus 1.5 × the interquartile range or less than the 25% quantile minus 1.5 × the interquartile range. **b)** Examples of predicted inter-species recombination in 6 vaccine candidates which were highly prevalent (>99%) in both SDSE and *S. pyogenes*. Species is indicated by colour: SDSE blue, *S. pyogenes* red, with donor species in a darker and recipient species in a lighter shade. Percent identity to reference *S. pyogenes* SF370 (NC_002737.2) is given at the top and reference SDSE GGS_124 (NC_012891.1) at the bottom of each plot over a 10bp sliding window. Recombination breakpoints predicted by fastGEAR given by vertical bars. *oppA* and *ropA* (TF) were predicted to be whole or near whole gene recombinants.

**Supplementary Table 1. a)** *Streptococcus dysgalactiae* subsp. *equisimilis* isolates included in this study with associated metadata. **b)** Virulence factor, antimicrobial resistance, and putative regulator screen. Putative stand-alone and two-component regulators were defined as by Buckley *et al.*[78, 79]

*See separate excel file*

**Supplementary Table 2. a)** List of 837 *Streptococcus dysgalactiae* subsp. *equisimilis* core genes identified by fastGEAR as having recombinogenic signatures. fastGEAR infers "recent" events as affecting individual isolates in the dataset whereas "ancestral" events affect an entire cluster of sequences. GGS_124 RefSeq locus tags (NC_012891.1) are given where available. Where not available, NC_019042.1 or NC_018712.1 locus tags are given except for 2 genes which were missing in all three RefSeq reference genomes. A small number of genes were merged in the pangenome by Panaroo[69] and locus tags are delimited by ";". A small number of genes were "refound" where additional annotations are inferred by Panaroo through comparison across the pangenome. **b)** List of 1,116 genes core in both *Streptococcus dysgalactiae* subsp. *equisimilis* (SDSE) and *S. pyogenes* analysed for putative cross-species recombination events by fastGEAR[35]. Core genes present in >99% of SDSE and *S. pyogenes* isolates with less than 25% length variation were included in the analysis. Of 1,166 core genes, 526 had at least two sequence clusters inferred by fastGEAR and at least one of which had members from both species (shared clusters). These may reflect whole gene recombination or shared ancestry. Furthermore, 393 genes with unique reference genome GGS_124 (NC_012891.1) SDSE locus tags had predicted recombination from a single-species sequence cluster to a genome from the other species (cross species events). For genes where a GGS_124 locus tag was not available, the locus tag from SDSE reference genome AC-2713 (NC_019042.1) was given.

*See separate excel file*

**Supplementary Table 3.** Ratio of SNPs due to recombination or mutation (r/m) for the 12 most sampled *Streptococcus dysgalactiae* subsp. *equisimilis* whole genome clusters as inferred by Gubbins[31]. Genomes from each genome cluster were aligned against an internal reference. For clusters with no complete genomes, genomes were aligned to the genome within the cluster with the highest N50.

| Whole genome cluster | No. isolates | Reference | Reference N50 | Reference contigs >1000bp | r/m (median, range) | No. recombination blocks/genome (median, range) | Length recombination segment (median, range) | No. recombination events (median, range) |
|---|---|---|---|---|---|---|---|---|
| 1 | 67 | SRR3676046 | 141405 | 39 | 5.48 (4.33, 10.12) | 43 (33, 80) | 5358 (45, 46903) | 43 (33, 80) |
| 2 | 59 | NC_018712.1 | - | 1 | 6.25 (2.86, 16.19) | 15 (8, 38) | 4936 (12, 55653) | 15 (8, 38) |
| 3 | 42 | SRR3676037 | 77623 | 52 | 7.05 (3.83, 10.19) | 64 (37, 88) | 5172 (6, 53277) | 64 (37, 88) |
| 4 | 41 | SRR3676051 | 136274 | 44 | 6.49 (4.14, 10.84) | 51 (41, 63) | 3810 (39, 43594) | 51 (41, 63) |
| 5 | 39 | NS3396 | - | 1 | 5.51 (0.34, 14.89) | 24 (2, 44) | 4062 (20, 41835) | 24 (2, 44) |
| 6 | 28 | NC_019042.1 | - | 1 | 3.48 (2.04, 4.64) | 50 (34, 59) | 4504 (28, 97789) | 50 (34, 59) |
| 7 | 28 | SRR3676017 | 105119 | 33 | 3.85 (2.12, 4.47) | 41 (29, 49) | 4884 (9, 40002) | 41 (29, 49) |
| 8 | 20 | SRR5754605 | 146734 | 39 | 2.24 (1.62, 2.76) | 7 (6, 9) | 3005 (69, 41458) | 7 (6, 9) |
| 9 | 19 | SRR3676111 | 70528 | 49 | 1.83 (1.06, 10.82) | 4 (2, 12) | 8094 (273, 26036) | 4 (2, 12) |

| | | | | | | | | |
|---|---|---|---|---|---|---|---|---|
| 10 | 15 | ERR1883109 | 75136 | 43 | 4.87 (2.79, 7.05) | 44 (26, 52) | 4856 (45, 37411) | 44 (26, 52) |
| 11 | 14 | NC_012891.1 | - | 1 | 4.05 (3.32, 4.44) | 60 (38, 67) | 5621 (15, 30903) | 60 (38, 67) |
| 12 | 13 | SRR3676040 | 62851 | 59 | 0.38 (0.28, 4.30) | 3 (2, 21) | 5579 (189, 20548) | 3 (2, 21) |

**Supplementary Table 4.** **a)** Prophage, integrative conjugative element and mobility element *Streptococcus dysgalactiae* subsp. *equisimilis* insertion regions mapped to the GGS_124 reference genome (NC_012891.1). **b)** Alternative *Streptococcus dysgalactiae* subsp. *equisimilis* insertion regions with fully assembled elements reflective of likely genome rearrangements around mobile genetic elements. Insertion regions were mapped to the GGS_124 reference genome (NC_012891.1). **c)** *Streptococcus pyogenes* integrative conjugative element and prophage insertion regions mapped to the *S. pyogenes* reference genome SF370 (NC_002737.2). Homologous *Streptococcus dysgalactiae* subsp. *equisimilis* (SDSE) insertion region no. refers to insertion region numbers given in Figure 3a.

*See separate excel file*

**Supplementary Table 5.** *S. pyogenes* candidate vaccine antigens examined in this study for presence in SDSE. Adapted from Davies *et al.*[22] Sequence locus tags are provided from *S. pyogenes* reference genome MGAS5005 (NC_007297.2) or SF370 (NC_002737.2). 180bp N-terminal *emm* sequences obtained from Centers for Disease Control and Prevention *emm* database (https://www2.cdc.gov/vaccines/biotech/strepblast.asp).

| **Antigen** | **Full length** | **Small peptide** | **Genbank accession or locus tag** | **Reference** |
|---|---|---|---|---|
| Streptolysin O (SLO)[@#^&] | | | M5005_RS00920 | Bensi *et al.* 2012[80] |
| Arginine deiminase (Adi)[#] | | | M5005_RS06290 | Henningham *et al.* 2012[81] |
| C5a peptidase (ScpA)[+#^&] | | | M5005_RS08460 | Ji *et al.* 1997[82] |
| Adhesion and division protein (SpyAD)[+@^&] | | | M5005_RS01310 | Bensi *et al.* 2012[80] |
| Oligopeptide-binding protein (OppA)[+] | | | M5005_RS01400 | Reglinski *et al.* 2016[83] |
| Streptococcal hemoprotein receptor (Shr) | | | M5005_RS07570 | Huang *et al.* 2011[84] |
| Putative pullulanase (PulA)[+] | | | SPY_RS08195 | Reglinski *et al.* 2016[83] |
| Trigger factor (TF)[#] | | | M5005_RS07965 | Henningham *et al.* 2012[81] |
| Nucleoside-binding protein (Spy0942)[+] | | | SPY_RS05150 | Reglinski *et al.* 2016[83] |
| Hypothetical membrane associated protein (Spy0762)[+] | | | SPY_RS04325 | Reglinksi *et al.* 2016[83] |
| Serine esterase (Sse) | | | M5005_RS06980 | Liu *et al.* 2007[85] |
| Cell surface protein (Spy0651)[+] | | | SPY_RS03510 | Reglinski *et al.* 2016[83] |
| Group A carbohydrate (GAC operon)[@^] | | | M5005_RS03095 to M5005_RS03150 | Sabharwal *et al.* 2006[86] |
| Serine protease (SpyCEP)[@#&] | | | M5005_RS01880 | Zingaretti *et al.* 2010[87] |

| | | | | |
|---|---|---|---|---|
| Serine protease SpyCEP: S2 peptide (S2) | | | ABD72239.1 amino acid 205 to 224 | Pandey *et al.* 2016[88] |
| Cysteine protease (SpeB) | | | M5005_RS08545 | Kapur *et al.* 1994[89] |
| Streptococcal immunoglobulin-binding protein 35 (Sib35) | | | AB254157.1 | Okamoto *et al.* 2005[90] |
| Streptococcal pyrogenic exotoxin A (SpeA) | | | X03929.1 | Ulrich 2008[91] |
| Streptococcal pyrogenic exotoxin C (SpeC) | | | SPY_RS02955 | McCormick *et al.* 2000[92] |
| M1 protein (M1) | | | M5005_RS08475 | Fox *et al.* 1973[93] |
| M protein: N-terminal peptide (30-valent) | | | CDC | Dale *et al.* 2011[94] |
| M protein: C-terminal peptide (J8) | | | See reference | Batzloff *et al.* 2003[95] |
| M protein: C-terminal peptide (StreptInCor) | | | See reference | Guilherme *et al.* 2006[96] |
| M protein: C-terminal peptide (P*17) | | | | Nordström *et al.* 2017[97] |
| Streptococcal fibronectin binding protein I (SfbI) | | | X67947.1 | Guzman *et al.* 1999[98] |
| Serum opacity factor (SfbII/SOF) | | | X83303.1 | Courtney *et al.* 2003[99] |
| Fibronectin-binding protein 54 (Fbp54) | | | AAA57236.1 | Kawabata *et al.* 2001[100] |
| Fibronectin binding protein A (FbpA) | | | M5005_RS08455 | Ma *et al.* 2009[101] |
| Streptococcal protective antigen (Spa) | | | AF086813.2 | Dale *et al.* 1999[102] |
| Rib-like cell wall protein (R28) | | | AF091393.1 | Stalhammar-Carlemalm *et al.* 1999[103] |
| Sortase A (SrtA)[&] | | | AAK34025.1 | Fan *et al.* 2014[104] |
| Multivalent T antigen subdomains (Tee)[%] | | | See Reference | Loh *et al.* 2021[105] |

[@] Component of the Novartis (GSK) combination vaccine (Di Benedetto *et al.* 2020) [106]
[+] Component of the Spy7 combination vaccine (Reglinski *et al*. 2016) [83]
[#] Component of the Combo#5 combination vaccine (Rivera-Hernandez *et al.* 2016) [107]
^ Component of VAX-A1 combination vaccine (Gao *et al.* 2021) [108]
[&] Component of 5CP combination vaccine (Bi *et al.* 2019) [109]
[%] Component of TeeVax combination vaccine which contains three multivalent fusion proteins, TeeVax1 (6 Tee antigens), TeeVax2 (7 Tee antigens), and TeeVax3 (5 Tee antigens) (Loh *et al.* 2021) [105]

**Supplementary Table 6.** Theoretical coverage of SDSE and *S. pyogenes* by combination vaccines based on genomes included in this study.

| **Vaccine** | **Antigen** | ***S. pyogenes*** **(n = 2083)** | **SDSE (n = 501)** |
|---|---|---|---|
| 4-component (GSK) [106] | SLO | >99% | >99% |
| | SpyCEP | >99% | 0% |
| | SpyAD | >99% | >99% |
| | GAC | >99% | 6% |
| | Any antigen | >99% | >99% |
| Spy7[83] | Spy0651 | >99% | 29% |
| | Spy0765 | >99% | >99% |
| | Spy0942 | >99% | >99% |
| | PulA | >99% | >99% |
| | OppA | >99% | >99% |
| | SpyAD | >99% | >99% |
| | ScpA | 98% | >99% |
| | Any antigen | >99% | >99% |
| Combo #5[107] | TF | >99% | >99% |
| | ScpA | 98% | >99% |
| | SpyCEP | >99% | 0% |
| | ADI | >99% | >99% |
| | SLO | >99% | >99% |
| | Any antigen | >99% | >99% |
| StrepInCor[96] | B cell epitope | 22% | 0% |
| | T cell epitope | 11% | 8% |
| | Common epitope | 18% | 74% |
| | Any epitope | 23% | 74% |

| | | | |
|---|---|---|---|
| S2.0 – J8.0[88] | S2.0 | 55% | 0% |
| | J8 | 37% | >99% |
| | Any epitope | 72% | >99% |
| 30-valent M[94] | 30 *emm* families | 48% | 1% |
| 30-valent with Mrp[110] | 30 *emm* families | 48% | 1% |
| | MrpI | 8% | 0% |
| | MrpII | 9% | 0% |
| | MrpIII | 5% | 0% |
| | Any antigen | 60% | 1% |
| VAX-A1[108] | GAC | >99% | 6% |
| | SpyAD | >99% | >99% |
| | ScpA | 98% | >99% |
| | SLO | >99% | >99% |
| | Any antigen | >99% | >99% |
| 5CP[109] | SrtA | >99% | >99% |
| | SpyAD | >99% | >99% |
| | ScpA | 98% | >99% |
| | SLO | >99% | >99% |
| | SpyCEP | >99% | 0% |
| | Any antigen | >99% | >99% |
| TeeVax[105] | TeeVax1 | 44% | 0% |
| | TeeVax2 | 40% | 6% |
| | TeeVax3 | 12% | 7% |
| | Any antigen | 85% | 13% |
| P*17 – S2.0[111] | P*17 | 37% | 87% |
| | S2.0 | 55% | 0% |
| | Any antigen | 72% | 87% |

**Supplementary references**


78. Buckley SJ, Davies MR, McMillan DJ. In silico characterisation of stand-alone response regulators of *Streptococcus pyogenes*. *PLoS One* 2020; **15**: e0240834.

79. Buckley SJ, Timms P, Davies MR et al. In silico characterisation of the two-component system regulators of *Streptococcus pyogenes*. *PLoS One* 2018; **13**: e0199163.

80. Bensi G, Mora M, Tuscano G et al. Multi high-throughput approach for highly selective identification of vaccine candidates: the Group A *Streptococcus* case. *Mol Cell Proteomics* 2012; **11**: M111.015693.

81. Henningham A, Chiarot E, Gillen CM et al. Conserved anchorless surface proteins as group A streptococcal vaccine candidates. *J Mol Med (Berl)* 2012; **90**: 1197-207.

82. Ji Y, Carlson B, Kondagunta A et al. Intranasal immunization with C5a peptidase prevents nasopharyngeal colonization of mice by the group A *Streptococcus*. *Infect Immun* 1997; **65**: 2080-7.

83. Reglinski M, Lynskey NN, Choi YJ et al. Development of a multicomponent vaccine for *Streptococcus pyogenes* based on the antigenic targets of IVIG. *J Infect* 2016; **72**: 450-9.

84. Huang YS, Fisher M, Nasrawi Z et al. Defense from the Group A *Streptococcus* by active and passive vaccination with the streptococcal hemoprotein receptor. *J Infect Dis* 2011; **203**: 1595-601.

85. Liu M, Zhu H, Zhang J et al. Active and passive immunizations with the streptococcal esterase Sse protect mice against subcutaneous infection with group A streptococci. *Infect Immun* 2007; **75**: 3651-7.

86. Sabharwal H, Michon F, Nelson D et al. Group A streptococcus (GAS) carbohydrate as an immunogen for protection against GAS infection. *J Infect Dis* 2006; **193**: 129-35.

87. Zingaretti C, Falugi F, Nardi-Dei V et al. *Streptococcus pyogenes* SpyCEP: a chemokine-inactivating protease with unique structural and biochemical features. *Faseb j* 2010; **24**: 2839-48.

88. Pandey M, Mortensen R, Calcutt A et al. Combinatorial Synthetic Peptide Vaccine Strategy Protects against Hypervirulent CovR/S Mutant Streptococci. *J Immunol* 2016; **196**: 3364-74.

89. Kapur V, Maffei JT, Greer RS et al. Vaccination with streptococcal extracellular cysteine protease (interleukin-1 beta convertase) protects mice against challenge with heterologous group A streptococci. *Microb Pathog* 1994; **16**: 443-50.

90. Okamoto S, Tamura Y, Terao Y et al. Systemic immunization with streptococcal immunoglobulin-binding protein Sib 35 induces protective immunity against group A *Streptococcus* challenge in mice. *Vaccine* 2005; **23**: 4852-9.

91. Ulrich RG. Vaccine based on a ubiquitous cysteinyl protease and streptococcal pyrogenic exotoxin A protects against *Streptococcus pyogenes* sepsis and toxic shock. *J Immune Based Ther Vaccines* 2008; **6**: 8.

92. McCormick JK, Tripp TJ, Olmsted SB et al. Development of streptococcal pyrogenic exotoxin C vaccine toxoids that are protective in the rabbit model of toxic shock syndrome. *J Immunol* 2000; **165**: 2306-12.

93. Fox EN, Waldman RH, Wittner MK et al. Protective study with a group A streptococcal M protein vaccine. Infectivity challenge of human volunteers. *J Clin Invest* 1973; **52**: 1885-92.

94. Dale JB, Penfound TA, Chiang EY et al. New 30-valent M protein-based vaccine evokes cross-opsonic antibodies against non-vaccine serotypes of group A streptococci. *Vaccine* 2011; **29**: 8175-8.

95. Batzloff MR, Hayman WA, Davies MR et al. Protection against group A *Streptococcus* by immunization with J8-diphtheria toxoid: contribution of J8- and diphtheria toxoid-specific antibodies to protection. *J Infect Dis* 2003; **187**: 1598-608.

96. Guilherme L, Faé KC, Higa F et al. Towards a vaccine against rheumatic fever. *Clin Dev Immunol* 2006; **13**: 125-32.

97. Nordström T, Pandey M, Calcutt A et al. Enhancing Vaccine Efficacy by Engineering a Complex Synthetic Peptide To Become a Super Immunogen. *J Immunol* 2017; **199**: 2794-802.

98. Guzmán CA, Talay SR, Molinari G et al. Protective immune response against *Streptococcus pyogenes* in mice after intranasal vaccination with the fibronectin-binding protein SfbI. *J Infect Dis* 1999; **179**: 901-6.

99. Courtney HS, Hasty DL, Dale JB. Serum opacity factor (SOF) of *Streptococcus pyogenes* evokes antibodies that opsonize homologous and heterologous SOF-positive serotypes of group A streptococci. *Infect Immun* 2003; **71**: 5097-103.
100. Kawabata S, Kunitomo E, Terao Y et al. Systemic and mucosal immunizations with fibronectin-binding protein FBP54 induce protective immune responses against *Streptococcus pyogenes* challenge in mice. *Infect Immun* 2001; **69**: 924-30.
101. Ma CQ, Li CH, Wang XR et al. Similar ability of FbaA with M protein to elicit protective immunity against group A *Streptococcus* challenge in mice. *Cell Mol Immunol* 2009; **6**: 73-7.
102. Dale JB, Chiang EY, Liu S et al. New protective antigen of group A streptococci. *J Clin Invest* 1999; **103**: 1261-8.
103. Stålhammar-Carlemalm M, Areschoug T, Larsson C et al. The R28 protein of *Streptococcus pyogenes* is related to several group B streptococcal surface proteins, confers protective immunity and promotes binding to human epithelial cells. *Mol Microbiol* 1999; **33**: 208-19.
104. Fan X, Wang X, Li N et al. Sortase A induces Th17-mediated and antibody-independent immunity to heterologous serotypes of group A streptococci. *PLoS One* 2014; **9**: e107638.
105. Loh JMS, Rivera-Hernandez T, McGregor R et al. A multivalent T-antigen-based vaccine for Group A *Streptococcus*. *Sci Rep* 2021; **11**: 4353.
106. Di Benedetto R, Mancini F, Carducci M et al. Rational Design of a Glycoconjugate Vaccine against Group A *Streptococcus*. *Int J Mol Sci* 2020; **21**.
107. Rivera-Hernandez T, Pandey M, Henningham A et al. Differing Efficacies of Lead Group A Streptococcal Vaccine Candidates and Full-Length M Protein in Cutaneous and Invasive Disease Models. *mBio* 2016; **7**.
108. Gao NJ, Uchiyama S, Pill L et al. Site-Specific Conjugation of Cell Wall Polyrhamnose to Protein SpyAD Envisioning a Safe Universal Group A Streptococcal Vaccine. *Infectious Microbes & Diseases* 2021; **3**: 87-100.
109. Bi S, Xu M, Zhou Y et al. A Multicomponent Vaccine Provides Immunity against Local and Systemic Infections by Group A *Streptococcus* across Serotypes. *mBio* 2019; **10**.

110. Courtney HS, Niedermeyer SE, Penfound TA et al. Trivalent M-related protein as a component of next generation group A streptococcal vaccines. *Clin Exp Vaccine Res* 2017; **6**: 45-9.
111. Ozberk V, Reynolds S, Huo Y et al. Prime-Pull Immunization with a Bivalent M-Protein and Spy-CEP Peptide Vaccine Adjuvanted with CAF®01 Liposomes Induces Both Mucosal and Peripheral Protection from covR/S Mutant *Streptococcus pyogenes*. *mBio* 2021; **12**.
